# High-resolution chromosome-level genome provides molecular insights into adaptive evolution in crabs

**DOI:** 10.1101/2024.06.24.600346

**Authors:** Yin Zhang, Ye Yuan, Mengqian Zhang, Xiaoyan Yu, Bixun Qiu, Fangchun Wu, Douglas R. Tocher, Jiajia Zhang, Shaopan Ye, Wenxiao Cui, Jonathan Y. S. Leung, Mhd Ikhwanuddin, Waqas Waqas, Tariq Dildar, Hongyu Ma

**Affiliations:** Guangdong Provincial Key Laboratory of Marine Biotechnology, Shantou University, Shantou, China; International Joint Research Center for the Development and Utilization of Important Mariculture Varieties Surrounding the South China Sea Region, Shantou University, Shantou, China; STU-UMT Joint Shellfish Research Laboratory, Shantou University, Shantou, China; Higher Institute Centre of Excellence (HICoE), Institute of Tropical Aquaculture and Fisheries, Universiti Malaysia Terengganu, Kuala Nerus, Terengganu, Malaysia

**Keywords:** whole genome, crab, early development, ovary maturation, LC-PUFA

## Abstract

Crabs thrive in diverse ecosystems, from coral reefs to hydrothermal vents and terrestrial habitats. Here, we report a comprehensive genomic analysis of the mud crab using ultralong sequencing technologies, achieving a high-quality chromosome-level assembly. The refined 1.21 Gb genome, with an impressive contig N50 of 11.45 Mb, offers a valuable genomic resource. Gene family analysis shows expansion in development-related pathways and contraction in metabolic pathways, indicating niche adaptations. Notably, Investigation into Hox gene regulation sheds light on their role in pleopod development, with the *Abd-A* gene identified as a linchpin. Posttranscriptional regulation involving novel-miR1317 negatively regulates *Abd-A* levels. Furthermore, the *fru* gene’s potential role in ovarian development and the identification of novel-miRNA-35 as a regulator of *Spfru2* add complexity to gene regulatory networks. Comparative functional analysis across Decapoda species reveals neofunctionalization of the *elovl6* gene in the synthesis of long-chain polyunsaturated fatty acids (LC-PUFA), suggesting its importance in environmental adaptation. These findings contribute significantly to our understanding of crab adaptability and evolutionary dynamics, offering a robust foundation for future investigations.

## Introduction

Evolutionary adaptation involves organisms adjusting to their environment to increase survival chances, resulting in diverse modifications including changes to body plans, development, and growth patterns. These adaptations are particularly evident among arthropods, specifically Decapoda crustaceans, which include economically valuable species such as shrimp, crayfish, lobsters, and crabs. Decapods occupy distinct ecological niches and employ diverse adaptive strategies, with shrimp navigating near the seafloor and crabs inhabiting burrows or crevices within muddy substrates during day. True crabs belong to the Decapoda infraorder Brachyura and can be traced back to the early mid-Jurassic period (Schweitzer and Feldmann 2010; Ma et al, 2019). The success of brachyuran crabs suggests that acquiring a crab-like morphology may have served as a pivotal innovation and, thus, they are an ideal group for studying trends in biodiversity over time (Morrison et al, 2002; Wolfe et al, 2021). Therefore, crabs provide an illuminating model system for investigating these foundations within Decapoda.

The majority of animals have intricate life cycles, which include larval and adult stages with distinct morphologies and ecological niches. In Decapoda, particularly in crabs, the morphological diversity is intriguing (Wolfe et al, 2019; Wolfe et al, 2021). Metamorphosis enables crabs to develop unique body shapes, and to transition from an oceanic planktonic to a benthic niche (Keiler et al, 2017). Crab larvae undergo metamorphosis from a laterally compressed zoea to a dorsoventrally compressed megalopa with crab-like features, recapitulating the morphological evolution of Decapoda. However, the evolutionary and genetic mechanisms underpinning crab metamorphosis remain inadequately understood (McNamara and Faria 2012; Bracken-Grissom et al, 2013; Scholtz 2014), despite indications of a correlation between body form and ecology (Luque et al, 2019). Concurrently, comprehending reproductive strategies is pivotal for elucidating evolutionary history, given the contribution these strategies make to enhancing parental fitness (Stearns 1992). While mammals exemplify the apogee of animal evolution with complex reproductive behaviors and sophisticated physiological mechanisms (Gong et al, 2023), in fish, gametes mature in the gonad before mating occurs (Turner et al, 2022). Similarly, shrimps mate after gonadal maturation (Zhang et al, 2019), whereas crabs mate after females undergo reproductive molting when their ovaries are immature, thus triggering ovary maturation.

In the domains of reproduction, metamorphosis, growth, and life history, fatty acids, specifically long-chain polyunsaturated fatty acids (LC-PUFA), fulfill essential functions (Tocher 2010; Ting et al, 2020; Závorka et al, 2023). The allocation of capacities in animals entails a delicate equilibrium between acquiring diverse fatty acids from diet and synthesizing LC-PUFA through multiple pathways regulated by fatty acid desaturase (*fads*) and elongase (*elovl*) genes (Twining et al, 2021; Castro et al, 2016). Initially, it was postulated that marine animals had limited ability to synthesize LC-PUFA due to the high availability of these fatty acids in marine ecosystems (Kabeya et al, 2018). However, recent reports indicate that euryhaline herbivorous fish and some invertebrates can also produce LC-PUFA endogenously (Monroig et al, 2018; Xie et al, 2021). Interestingly, the genes encoding different types of Elovl are more diverse in invertebrates than vertebrates (Kabeya et al, 2018; Monroig et al, 2022; Ramos-Llorens et al, 2023), although this diversity remains poorly understood.

The utilization of genome sequencing is crucial for understanding organism evolution and identifying genes involved in animal adaptations (Zhang et al, 2019; Yim et al, 2014; Gutekunst et al, 2018). Despite the availability of several genome maps for shrimps and crabs (Zhang et al, 2019; Song et al, 2016; Tang et al, 2021; Tang et al, 2020; Zhao et al, 2021; Cui et al, 2021; Chebbi et al, 2019), the current genomic data for Decapoda are insufficient to facilitate comprehensive studies on development, reproduction, and nutrient acquisition within this ancient, successful lineage. Attaining a high-quality genome is imperative for establishing an experimental model to understand the molecular mechanisms underlying adaptive evolution in crabs. The current study presents a comprehensive analysis of the genome of the green mud crab *Scylla paramamosain*, providing a robust model system to investigate the molecular foundations of adaptive evolution within Decapoda.

## Results

### Genome sequencing, assembly and annotation

A total of 98.60 Gb pass reads were obtained using Nanopore ultralong-read sequencing. After quality control, genome assembly and correction, the *S. paramamosain* (Fig. 1A) genome had a size of 1.21 Gb with a contig N50 of 11.45 Mb. In total, 421 contigs with an average length of 11,740,790 bp were obtained and further assembled into 233 scaffolds with a scaffold N50 of 23.61 Mb (Table 1), which was longer than those of other crustaceans (Zhang et al, 2019; Gutekunst et al, 2018; Song et al, 2016; Tang et al, 2021; Cui et al, 2021; Chebbi et al, 2019) (Table S1). The N50 of the sequences obtained here is approximately five times that achieved in the previous genome assembly (N50 < 200 kb) (Zhao et al, 2021), indicating a significant improvement compared to the previous *S. paramamosain* genome. Overall, 95.26% of the 1,013 Arthropoda Benchmarking Universal Single-Copy Orthologs (BUSCOs) were identified in the assembled genome by searching against Arthropoda_odb 10. The contigs were further anchored to 49 pseudochromosomes with a total length of 1,106,215,978 bp, representing 99.47% of the genome (Fig. 1B). Chromosome 01 (chr01) is the largest, with a length of 42.18 Mb, whereas chr49 has the shortest length (5.67 Mb). In addition, chr06 has a strongly sex-linked signal using genome-wide association studies (GWAS) (Fig. 1C and Fig. S1), which have been conducted to identify genes associated with various economically relevant traits, including sex, growth, disease resistance and environmental adaptation, in shrimp (Zhang et al, 2019; Lyu et al, 2021). Previously, we identified 13 sex-specific SNPs (Waiho et al, 2019) located on chr06 in *S. paramamosain*, which indicated that chr06 is the sex chromosome of green mud crab. We observed a peak of Ks ranging from 0 to 1 and a summit at 0.2, representing a potential ancient whole-genome duplication (WGD) (Fig. 1D).

**Fig. 1.**
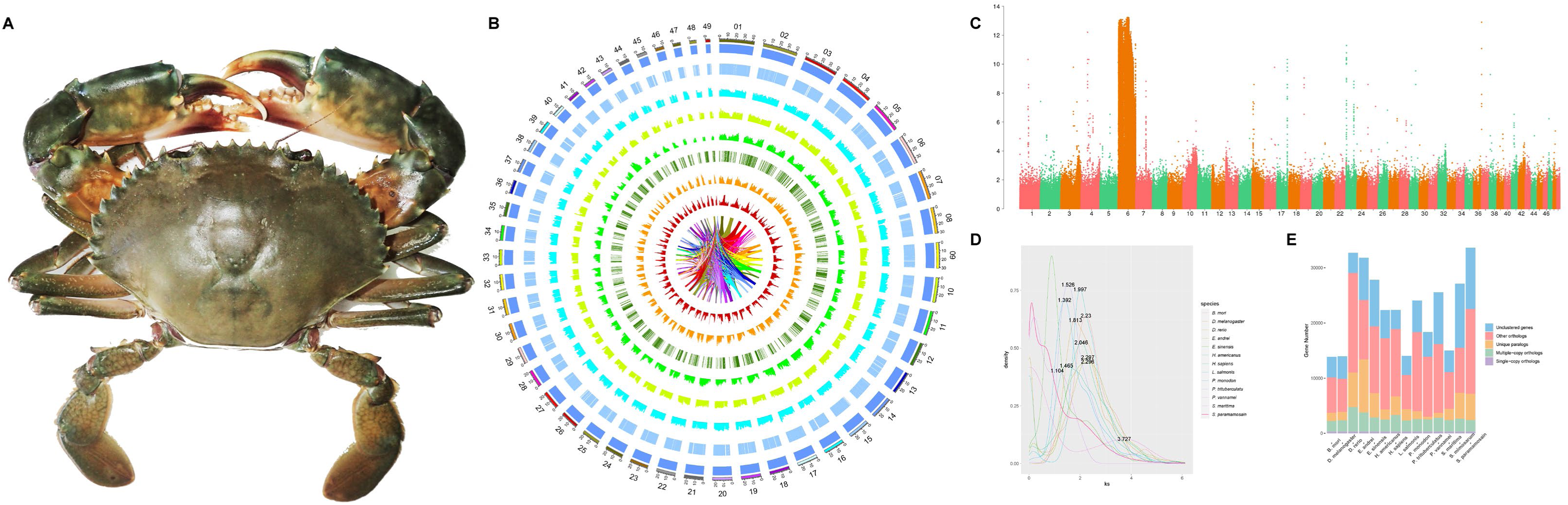
Schematic representation of the genomic characteristics of mud crab *Scylla paramamosain*. **(A)** Appearance of *S. paramamosain*. **(B)** Schematic representation of the genomic characteristics of *S. paramamosain*. From the outer to inner, Track 1: The 49 chromosomes of *S. paramamosain*; Track 2: Scaffolds anchored to each chromosome; Track 3: Protein-coding genes; Track 4: Distribution of gene density across the genome; Track 5: Distribution of GC content in the genome; Track 6: Distribution of 5 significantly expanded gene families in the genome; Track 7: Distribution of SSRs in the genome; Track 8: Distribution of transposable elements in the genome; Track 9: Schematic presentation of major interchromosomal relationships in the mud crab. **(C)** Sex linked region in the chromosomes of *S. paramamosain*. **(D)** Whole genome duplicates event in 12 species including *S. paramamosain, Portunus trituberculatus, Eriocheir sinensis, Penaeus vannamei, Homarus americanus, Penaeus monodon, Drosophila melanogaster, Bombyx mori, Eisenia andrei, Stegodyphus mimosarum, Strigamia maritima, Danio rerio*. **(E)** Single-copy orthologs are defined as orthologs that were present as a single-copy gene in all 14 species (*S. paramamosain, P. trituberculatus, E. sinensis, P. vannamei, H. americanus, P. monodon, D. melanogaster, B. mori, E. andrei, Lepeophtheirus salmonis, S. mimosarum, S. maritima, D. rerio,* and *Homo sapiens*). Multiple-copy orthologs represent the gene groups present in all species with a gene number of > 1 in at least one species. Species-specific paralogs represent genes uniquely present in only one species. Other types of orthologs represent the gene groups that are absent in some species and not species-specific paralogs.

**Table 1.** Summary of *Scylla paramamosain* genome assembly

| Stat Type | Contig Length<br>(bp) | Contig<br>Number | Scaffold Length<br>(bp) | Scaffold<br>Number | Gap<br>Length<br>(bp) | Gap<br>Number |
| --- | --- | --- | --- | --- | --- | --- |
| N50 | 11,448,802 | 37 | 23,607,973 | 20 | 100 | 113 |
| N60 | 9,208,292 | 49 | 21,816,234 | 25 | 100 | 135 |
| N70 | 6,865,899 | 64 | 19,336,055 | 31 | 100 | 158 |
| N80 | 3,900,000 | 87 | 15,570,585 | 38 | 100 | 180 |
| N90 | 1,000,000 | 151 | 9,316,856 | 47 | 100 | 203 |
| Longest | 30,957,417 | 1 | 42,181,690 | 1 | 100 | 225 |
| Total | 1,210,737,920 | 458 | 1,210,760,420 | 233 | 22,500 | 225 |
| Length>=1kb | 1,210,737,920 | 458 | 1,210,760,420 | 233 | 0 | 0 |
| Length>=2kb | 1,210,737,920 | 458 | 1,210,760,420 | 233 | 0 | 0 |
| Length>=5kb | 1,210,737,920 | 458 | 1,210,760,420 | 233 | 0 | 0 |

The assembly of the genome can be affected by various factors, such as individual heterozygosity, repetitive sequences, and GC bias (Zhang et al, 2012; Li et al, 2017). In addition, long reads can span complex or repetitive regions with a single continuous read, thus eliminating ambiguity in the positions or size, contributing to a more complete genome assembly (Jain et al, 2018). Herein, we performed ultralong-read sequencing and detected 587.19 Mb of repetitive sequences, accounting for 58.97% of the genome, which is much higher than that in Chinese mitten crab *Eriocheir sinensis* (Cui et al, 2021) and Pacific white shrimp *Litopenaeus vannamei* (Zhang et al, 2019), but lower than that in other crustaceans for which genome data are available (Gutekunst et al, 2018; Zhao et al, 2021). The genome contains 206,695 simple sequence repeats (SSRs), representing 0.25% of the total genome. Dinucleotide repeats were the dominant type of SSRs, accounting for 58.85% of the total SSRs. Moreover, there were 345,610 tandem repeats (TRs) with a length of 24,183,525 bp, representing 2.00% of the whole genome, which is much lower than that in Chinese mitten crab (Cui et al, 2021). As a predominant force driving genome expansion and evolution (Bire and Rouleux-Bonnin 2012; Elliott and Gregory 2015), transposable elements (TEs), with a proportion of 48.50% of the *S. paramamosain* genome, contain 17.40% long interspersed elements (LINEs), 8.22% long terminal repeats (LTRs), 22.05% DNA transposons, 0.33% rolling circle (RC) and 0.35% miniature inverted-repeat transposable elements (MITEs) (Fig. S2 and Table S2).

In *S. paramamosain*, 33,662 protein-coding genes were predicted with an average gene length of 12,535.06 bp, CDS length of 1,208.52 bp, exon length of 225.29 bp, and intron length of 2,595.33 bp (Table S3 and Fig. S3). Compared with Chinese mitten crab *E. sinensis* (Cui et al, 2021) and marbled crayfish *Procambarus virginalis* (Gutekunst et al, 2018), we found that *S. paramamosain* had relatively more coding genes. The average CDS length of green mud crab protein-coding genes was similar to that of Chinese mitten crab *E. sinensis* (Song et al, 2016) and shrimp *L. vannamei* (Zhang et al, 2019), but longer than that of swimming crab *Portunus trituberculatus* (Tang et al, 2020) (Table S1). Comparisons of gene number and protein length indicated superior quality in the green mud crab gene models compared to those of the Chinese mitten crab and swimming crab, suggesting useful resources for further comparative studies and a more comprehensive understanding of genomic characteristics in green mud crab. In total, 92.31% of the genes were annotated to public databases, including SwissProt (14,738 genes), InterPro (28,920 genes), Kyoto Encyclopedia of Genes and Genomes (KEGG; 11,128 genes), Gene Ontology (GO; 12,328 genes), EuKaryotic Orthologous Groups (KOG; 12,381 genes) and Non-Redundant Protein Sequence Database (Nr, 20,927 genes) (Fig. S4). We also annotated noncoding RNAs, including 449 miRNAs, 1,055 rRNAs, 36 snRNAs, and 3,153 tRNAs (Table S4).

### Gene family expansion, contraction and genome evolution

Most expansion events are associated with the adaptive evolution of specific phenotypes. To conduct gene family expansion and contraction events in *S. paramamosain*, gene family clustering analysis was conducted on 14 species, including five other crustacean species (*P. trituberculatus, E. sinensis, Penaeus vannamei, Homarus americanus*, and *P. monodon*), six insect species (*Drosophila melanogaster, Bombyx mori, Eisenia andrei, Lepeophtheirus salmonis, Stegodyphus mimosarum*, and *Strigamia maritima*), and two vertebrate species (*Danio rerio* and *Homo sapiens*). The results of the gene family analysis were categorized and statistically analyzed for each category. *S. paramamosain* had 297 single-copy genes, 4,775 unique genes, and 11,097 unclustered genes. Furthermore, the unclustered and unique genes were combined to yield a total of 15,872 species-specific genes (Fig. 1E and Fig. S5). These species-specific genes were significantly enriched in aminoacyl-tRNA biosynthesis, TNF signaling pathway, RNA degradation, ubiquitin mediated proteolysis, longevity regulating pathway, circadian entrainment, autophagy–animal, neurotrophin signaling pathway, adrenergic signaling in cardiomyocytes, MAPK signaling pathway, NF-kappa B signaling pathway, mTOR signaling pathway and c-type lectin receptor signaling pathway (Fig. S6), most of which are related to the development and lifestyle of the green mud crab. Phylogenetic analysis based on 297 single-copy orthologous genes suggested that crabs diverged from shrimp (represented by *L. vannamei*) ∼481.54 million years ago (Mya), that marine crabs diverged from freshwater crabs (*E. sinensis*) ∼280.48 Mya, and that *S. paramamosain* diverged from *P. trituberculatus* ∼141.1 Mya (Fig. S7).

A total of 12,274 gene families were identified through family clustering, with 725 being specific to *S. paramamosain*. Of the 33,662 genes, 22,565 genes could be classified into distinct gene families, with an average of 1.84 genes per family. As variations in gene copy number might support adaptive evolution, we examined the expansion and contraction of gene families in the *S. paramamosain* genome to explore the potential mechanisms underlying the adaptability of crabs. The number of expanded gene families in *S. paramamosain* was 1,545, while the number of contracted families was 2,671 (Fig. 2). Among them, the expanded genes were significantly enriched in development-related pathways such as betalain biosynthesis, ubiquitin-mediated proteolysis, circadian entrainment, neurotrophin signaling pathway, adrenergic signaling in cardiomyocytes, longevity regulating pathway, MAPK signaling pathway and autophagy–animal. Interestingly, when comparing shrimps and crabs, the expanded gene families in *S. paramamosain* were also significantly associated with these pathways (Table S5), suggesting that the specific developmental pattern of green mud crabs may be attributed to the expansion of these gene families. On the other hand, the contracted gene families were mainly annotated to nutritional metabolism, including carbon metabolism, propanoate metabolism, glyoxylate and dicarboxylate metabolism, microbial metabolism in diverse environments, biosynthesis of secondary metabolites, carbon fixation pathways in prokaryotes, methane metabolism, pyruvate metabolism, and glycolysis/gluconeogenesis (Fig. S8), indicating that there have been alterations in nutrient intake during the process of evolution. Therefore, the aforementioned species-specific and expanded gene families may have been important to *S. paramamosain* lineage-specific environmental adaptations, enabling the crab to occupy diverse niches in a myriad of stressful conditions.

**Fig. 2.**
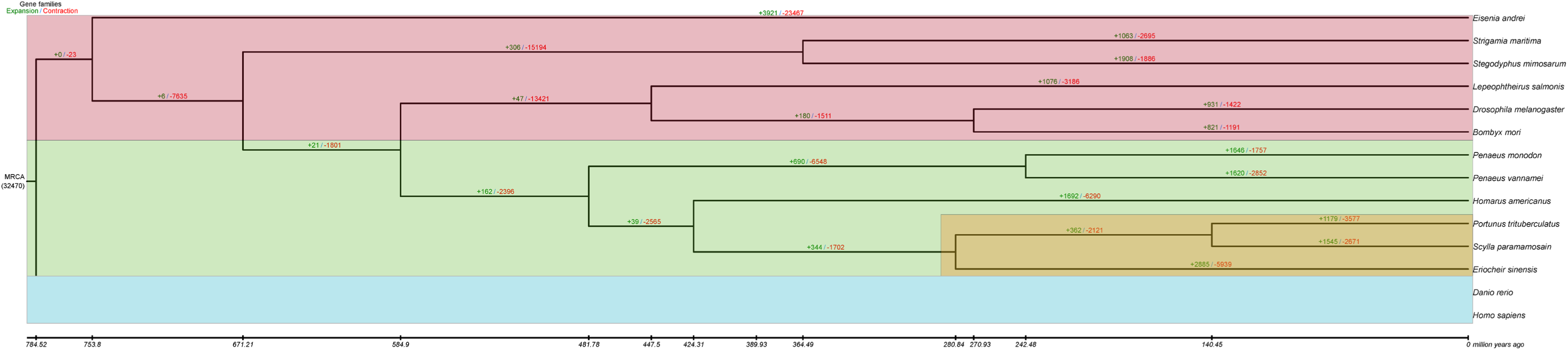
Phylogenetic tree and gene family expansion/contraction of *Scylla paramamosain.* Phylogenetic tree inferred from 338 single-copy orthologs among 14 selected species (*S. paramamosain, Portunus trituberculatus, Eriocheir sinensis, Penaeus vannamei, Homarus americanus, Penaeus monodon, Drosophila melanogaster, Bombyx mori, Eisenia andrei, Lepeophtheirus salmonis, Stegodyphus mimosarum, Strigamia maritima, Danio rerio,* and *Homo sapiens*). Numbers of expanded gene families are marked in green, and numbers of contracted gene families are marked in red. The number below the MRCA (most recent common ancestor) represents the total number of orthologs from OrthoMCL analysis used as input for CAFE expansion/contraction analysis.

### The function and regulatory mechanisms of Hox genes in the development of pleopods

Crustaceans present the most impressive diversity in body plan among arthropods (Deutsch and Mouchel-Vielh 2003). Hox genes encode homeodomain-containing transcription factors that play crucial roles in directing tissue differentiation and morphological development throughout all the principal axes of an embryo (Yan et al, 2019). Here, we found that the *S. paramamosain* genome contained all ten canonical Hox gene clusters (*lab, pb, Hox3, Dfd, Scr, ftz, Antp, Ubx, Abd-A*, and *Abd-B*) (Fig. 3 A, B), which were found in the arthropod ancestor (Grenier et al, 1997; Hughes and Kaufman 2002). The Hox gene clusters were located on the same chromosome (Fig. 3A), similar to the spatial collinearity in most bilaterians (Uengwetwanit et al, 2021). However, most Hox gene sequences contain breaks because of low-quality sequencing data, preventing the verification of the transcribed direction of the Hox gene cluster (Kim et al, 2018). Herein, the continuous location of the Hox gene cluster strongly supported the high integrity of our genome assembly of *S. paramamosain*. However, the expression of the Hox genes did not show whole-cluster temporal collinearity (Fig. 3C), similar to vertebrates, but exhibited subcluster-level temporal collinearity, similar to Chinese mitten crab *E. sinensis* (Cui et al, 2021), shrimp *P. monodon* (Uengwetwanit et al, 2021), scallop *Patinopecten yessoensis* (Wang et al, 2017), oyster *Crassostrea gigas* (Zhang et al, 2012) and sea squirt *Ciona intestinalis* (Ikuta et al, 2004).

**Fig. 3.**
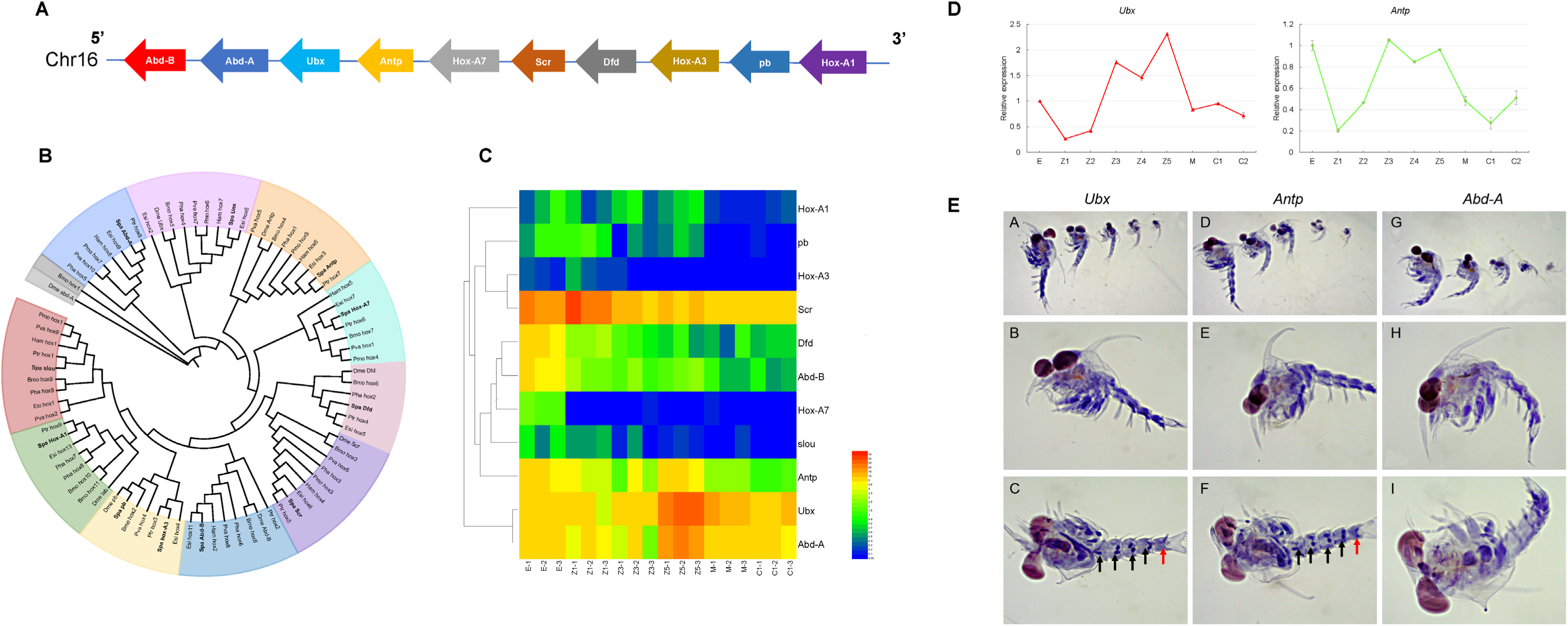
The conserved hox gene clusters and their expressions in *Scylla paramamosain*. **(A)** Location and transcript direction in the genome of *S. paramamosain* Hox genes. **(B)** Distribution and relationships of Hox gene cluster in *S. paramamosain*, *Bombyx mori* (Bmo), *Drosophila melanogaster* (Dme), *Eriocheir sinensis* (Esi), *Homarus americanus* (Ham), *Parhyale hawaiensis* (Pha), *Penaeus monodon* (Pmo), *Penaeus vannamei* (Pva), and *Portunus trituberculatus* (Ptr). **(C)** Temporal expression of *S. paramamosain* Hox cluster genes. E: embryo, Z1: zoea I, Z3: zoea III, Z5: zoea V, M: megalopa stage, and C1: crablet I. **(D)** Expressions of *Ubx* and *Antp* in early different developmental stages of *S. paramamosain*. **(E)** The expression locations of *Ubx*, *Antp* and *Abd-A* in *S. paramamosain* larvae.

Green mud crab larval includes stages from zoea I to megalopa, with zoea I has sessile eyes; zoea II has stalked eyes; zoea III has six segments on abdomen; zoea IV has the presence of pleopod buds and zoea V has the presence of setae on pleopod well developed and also the presence of setae. During the early stages of development in *S. paramamosain*, *SpUbx*, *SpAntp,* and *SpAbd-A* exhibited similar expression patterns that were upregulated during zoea stages III and V (Fig. 3C), suggesting their potential involvement in regulating zoeal development process. Additionally, there was a significant decrease in the expression levels of both *SpUbx* and *SpAntp* within the abdomen of zoea stage III to V individuals (Fig. 3D), while there was an increase in *SpAbd-A* (Fig. 4A). Given that green mud crab zoea stages III to V are critical periods for abdominal and maxillary development, it can be speculated that these three genes primarily function on the abdomen and maxillae of zoea larvae. *SpAbd-A* gene expression was significantly higher in the abdomen (Fig. 4A); however, at the protein level, it was higher in the cephalothorax (Fig. 4B). Furthermore, there was a gradual increase followed by a decrease in SpAbd-A protein expression in the abdomen of zoea I to zoea V, with a peak observed in the zoea stage III (Fig. 4B). The SpAbd-A protein was primarily expressed in maxillopods, pereiopods, and abdominal segments in the zoea (Fig. 4C). Interference with the expression level of *SpAbd-A* during the zoea stage IV resulted in the absence of pleopods in larvae (Fig. 4D), highlighting its crucial role in initiating and developing pleopods in *S. paramamosain* larvae. In crustaceans, *Abd-A* establishes segmental identity and boundaries; without *Abd-A*, present abdominal segments may develop thoracic-like limbs (Martin et al, 2016). The loss of *Abd-A* has also been associated with a lack of an abdomen in lineage Cirripedia (Mouchel-Vielh et al, 1998). Considering that morphological changes are mainly driven by alterations in gene expression patterns, it can be speculated that abdominal degeneration in crabs may be linked to low levels of *Abd-A* during brachyurization metamorphosis.

**Fig. 4.**
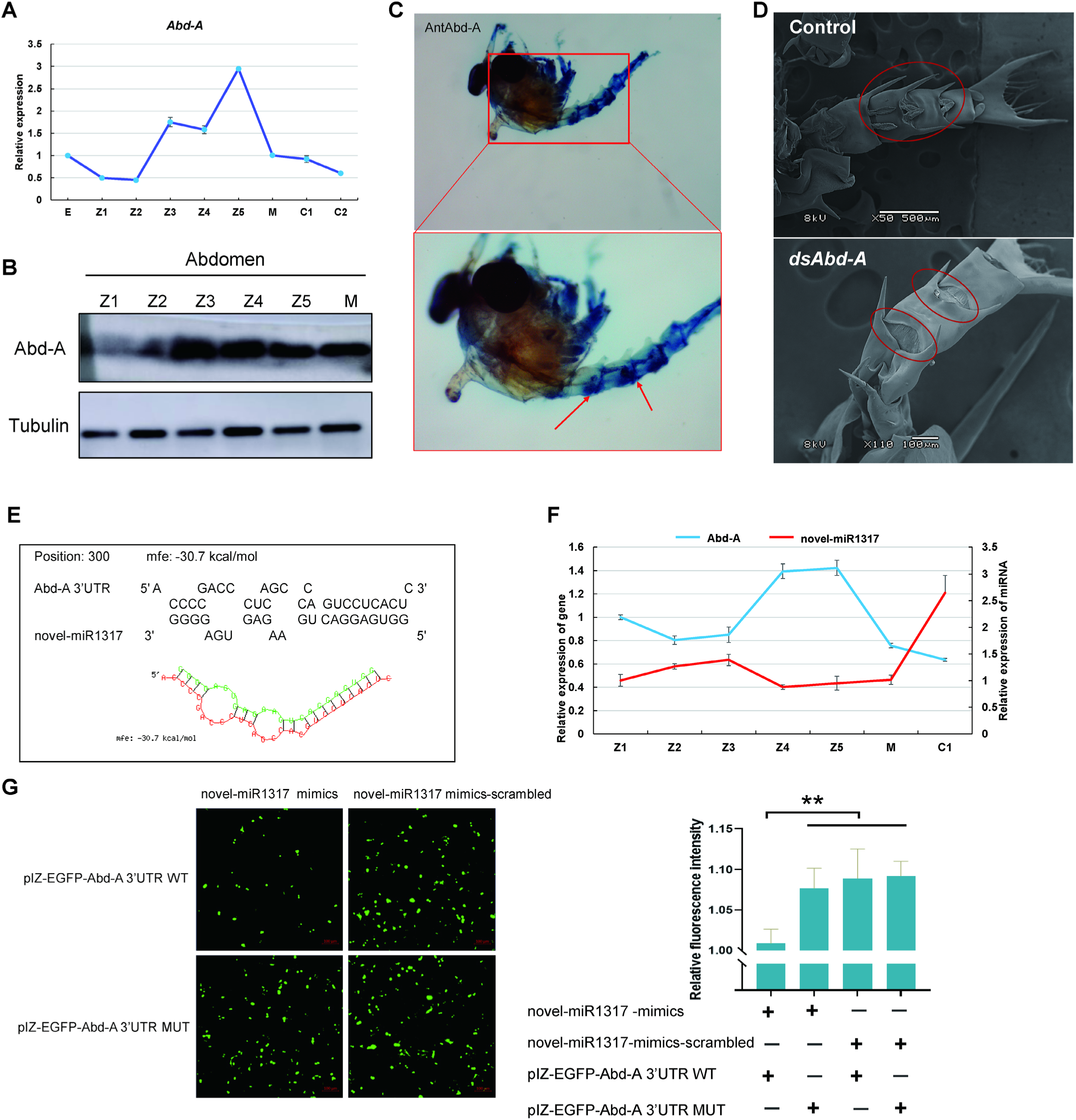
The function and regulation of Abd-A gene in *Scylla paramamosain* larvae. **(A)** Expressions of *Abd-A* in different stages of *S. paramamosain* larvae. **(B)** Expression of SpAbd-A protein in the abdomen of *S. paramamosain* larvae. **(C)** Expression locations of SpAbd-A protein in *S. paramamosain* larvae. **(D)** Phenotype changes of *S. paramamosain* at Zoeal stage V after RNAi at Zoeal stage IV. **(E)** Prediction binding sites of *Abd-A* and novel-miR1317. **(F)** Negative expressions patterns between novel-miR1317 and *Abd-A* in *S. paramamosain* larvae. **(G)** The S2 cells were co-transfected with WT Abd-A 3’-UTR, and the mutated-type of Abd-A 3’-UTR (MT), together with novel-miR1317 mimics or negative control mimic. The data were expressed as the relative fluorescence intensity. Z1-5: zoea I-V, M: megalopa stage, C1: crablet I.

Posttranscriptional regulation also fine-tunes Abd-A levels, wherein microRNAs such as miR-10 target *Abd-A* transcripts for degradation (Miura et al, 2011). The potential target miRNAs of *SpAbd-A* were identified through transcriptome data analysis of *S. paramamosain* larvae and RNAhybrid: novel-miR1317 was found to be one of them (Fig. 4E). A negative relationship between the expression level of novel-miR1317 and *SpAbd-A* was detected (Fig. 4F), verifying that novel-miR1317 negatively regulated the expression of the *Abd-A* gene, possibly facilitating the development of pleopods. In most insect species, miR-iab-4 exerts a negative regulatory effect on *Abd-A* and *Ubx* (Mouchel-Vielh et al, 1998), while miR-iab-8 regulates *Abd-A* and *Abd-B* (Garaulet and Lai 2015). In both Chinese mitten crab and swimming crab genomes, miR-993, miR-10, miR-iab-4, and miR-iab-8 were identified within the intergenic region of Hox gene clusters (Cui et al, 2021). Herein, the current study has demonstrated the significant role of *SpAbd-A* in the development of abdominal pleopods in *S. paramamosain* larvae. Its expression is negatively regulated by novel-miR1317 (Fig. 4G). These findings provide a theoretical foundation for further understanding the mechanisms of pleopod development of *S. paramamosain* larvae.

### Gene regulation of ovarian development

Reproductive molting is crucial for the reproductive process of crustaceans. After reproductive molting, female crabs mate with males, and the ovaries begin to develop rapidly (Li et al, 2022; Liu et al, 2020; Long et al, 2020; Wang et al, 2020). Numerous studies have demonstrated that the fruitless (*fru*) gene plays a role in the sex determination pathway in fly *D. melanogaster* (Manoli et al, 2005; Cachero et al, 2010). Meanwhile, we obtained the transcript fragments of fruitles2 (*Spfru2*) according to the gonadal transcriptome data of green mud crab (Yang et al, 2017). Additionally, we found that the expression levels of *Spfru2* in the gonad, thoracic ganglion, gill and muscle of females were significantly higher than those in males (Fig. S9A). Moreover, the expression level of *Spfru2* was the highest in ovarian stage III of the five different developmental stages (Fig. 5A). As the ovary increased in volume during stage III, the internal germ cells began to mature and were predominantly stage 3 oocytes (Wu et al, 2020). Additionally, *Spfru2* was mainly localized in the cytoplasm of oocytes, and, in males, it was detected mainly in the epithelia of seminiferous tubules and spermatids (Fig. 5B). This implies that it may function in ovarian maturation in green mud crab.

**Fig. 5.**
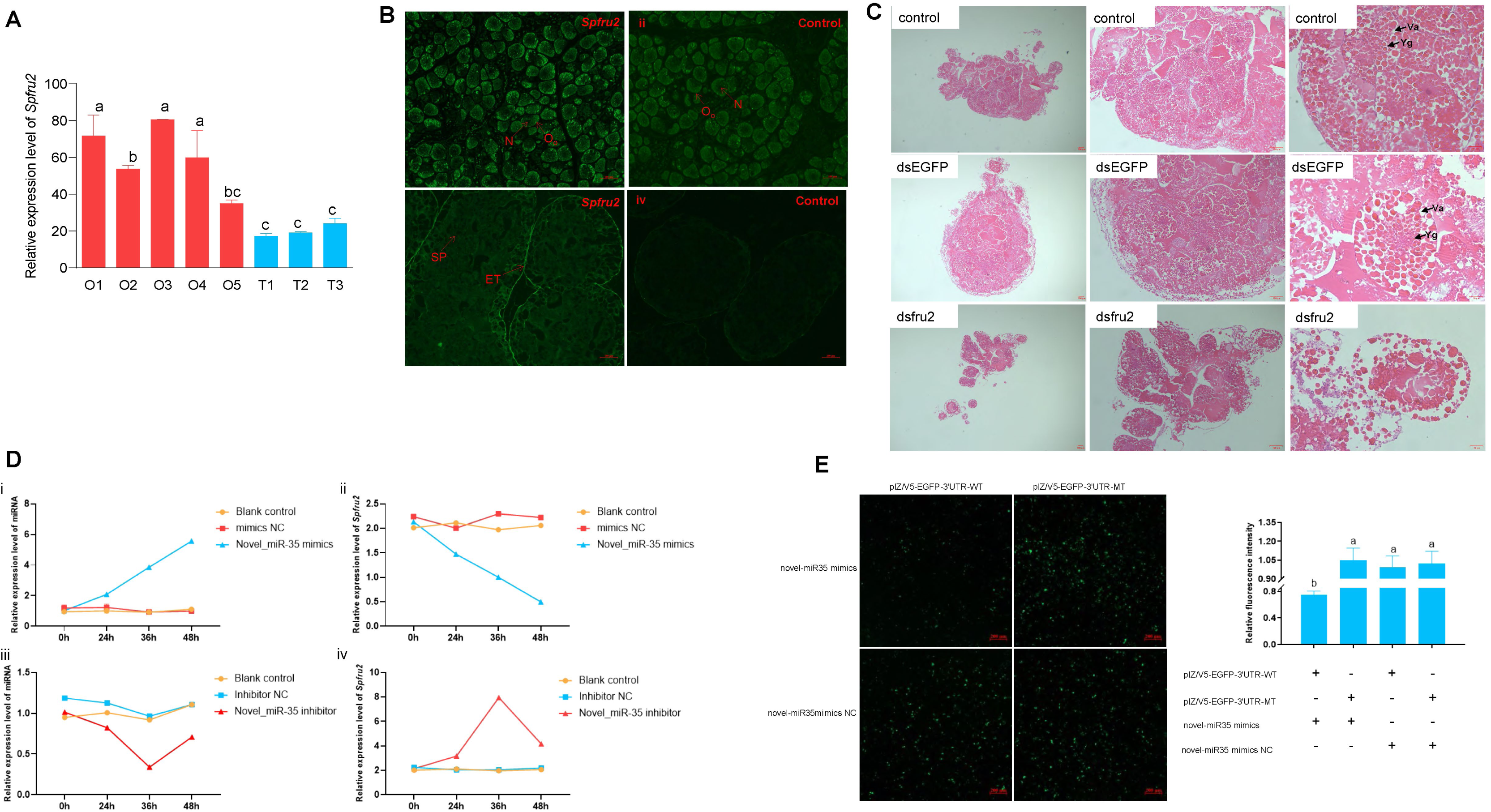
The expression and regulation of fru2 genes in gonads of *Scylla paramamosain*. **(A)** Relative expression of *Spfru2* in different developmental stages of the gonads. O: ovary; T: testis; O1-O5: stage OⅠ to stage OⅤ; T1-T3: stage TⅠ to stage TⅢ. Bars with different lowercase letters indicate significant difference (*p* < 0.05). **(B)** Distribution of *Spfru2* in ovary (i, ii) and testis (iii, iv) of *S. paramamosain*. N: nucleus; Oo: oocyte; ET: epithelia; SP: spermatid. Scale bars: 100 μm. **(C)** Phenotype changes of the ovary tissue after the interference of *fru2* expression. Yg: yolk granule; Va: Vacuole. **(D)** Relative expression of novel-miR35 and *Spfru2* of the ovary after overexpression and silencing of *novel-miR35. i: Relative expression of novel-miR35 in the ovary after overexpression of novel-miR35. ii: Relative expression of *Spfru2* in the ovary after overexpression of novel-miR35. iii: Relative expression of novel-miRNA-35 in the ovary after silencing of novel--miR35. iv: Relative expression of *Spfru2* in the ovary after silencing of novel-miR35. Bars with different lowercase letters indicate significant difference (*p* < 0.05). **(E)** The S2 cells were co-transfected with WT *fru2* 3’-UTR, and the mutated-type of *fru2* 3’-UTR (MT), together with novel-miR35 mimics or negative control mimic. The data were expressed as the relative fluorescence intensity.

The expression level of *Spfru2* was higher at day three post mating (Fig. S9B). In crabs, after the reproductive molt, females serve as the catalyst for the mating process and trigger ovarian development, including oogenesis (Yang et al, 2022). Oogenesis represents a metabolically demanding reproductive process that encompasses distinct phases. The subsequent stages of oogenesis involve primary and secondary vitellogenesis (Tsukimura 2001), which entail significant enlargement of oocyte diameter along with yolk protein accumulation within maturing oocytes. Subsequent to ovulation and release from the ovaries, eggs undergo fertilization through fusion with sperm residing within female spermathecae for varying durations ranging from one to several months (Tsukimura 2001). This observation further highlights the variable period required for sperm-egg maturation throughout the reproductive cycle of green mud crabs. The *in vitro* tissue culture experiments demonstrated a significant decrease in *Spfru2* expression at 8 h (Fig. S9C), indicating the effectiveness of RNA interference. Additionally, compared to the positive and negative controls, the treatment group that underwent *fru2* gene interference exhibited incomplete and fragmented egg cell morphology at 8 h (Fig. 5C). These findings suggest that the *fru2* gene also play a pivotal role in maintaining egg cell morphology, thereby influencing female gonadal development and maturation.

Furthermore, we found the expression of novel-miR35 and *Spfru2* in different gonad development stages displayed opposite expression patterns in both female and male crabs (Fig. S9). Within 48 h after injection of miRNA mimic/inhibitor, the expression levels of novel-miR35 and *Spfru2* had opposite variant trends in the gonads (Fig. 5D). We further validated the interaction between novel-miR35 and *Spfru2* using *in vitro* green fluorescent reporter assays. Both the novel-miR35 mimic and pre–novel-miR35 plasmid effectively reduced fluorescence intensity when co-transfected with the WT 3′-UTR reporter plasmid into *Drosophila* S2 cells, but this effect was largely restored for the co-transfected plasmid containing the mutant type (MT) 3′-UTR (Fig. 5E, F). These findings suggest that *Spfru2* might be a target of novel-miR35 in green mud crab.

### Neo-functionalization of the *elovl6* gene in the LC-PUFA synthesis pathway

The LC-PUFA are defined as fatty acids with ≥ 20 carbons and ≥ 3 double bonds and, depending upon the position of the last double bond relative to the methyl end, can be categorized into n-6 and n-3 series LC-PUFA, and are critical nutrients for animal reproduction and metamorphosis (Xie et al, 2021). Among them, n-3 LC-PUFA, especially eicosapentaenoic acid (EPA, 20:5n-3) and docosahexaenoic acid (DHA, 22:6n-3), contribute to organism health *via* multiple pathways and play crucial roles in cell membrane lipid functions, neurodevelopment, and immune responses, and have well-known beneficial effects in mitigating several diseases (Xie et al, 2021). Generally, LC-PUFA can be enriched in body tissues either by dietary acquisition or, in some animal species, through endogenous biosynthesis (Xie et al, 2021). The fatty acid elongase, Elovl6, as a crucial enzyme in the endogenous synthesis of fatty acids, participates in fatty acid metabolism, energy balance, insulin resistance, and metabolic diseases (Matsuzaka et al, 2007; Shimano 2012). To illuminate the neofunctionalization of the *elovl6* gene and its involvement in LC-PUFA biosynthesis in green mud crab, we first analyzed the molecular characteristics of the *Spelovl6* gene. The variable promoter of the *elovl6* gene in *S. paramamosain* resulted in the generation of three transcripts (*elovl6a*, *elovl6b*, and *elovl6c*) through splicing (Fig. 6A). Elovl6a, Elovl6b, and Elovl6c exhibited the typical structural features of Elovl protein family members (Fig. 6B). During embryonic development, the expression levels of the *elovl6* genes showed an overall increasing trend, reaching highest levels in the prehatching period, and decreasing significantly in the hatching period (Fig. S10 A, B, C). The expression levels of *elovl6* genes in the larval stages showed a generally stable trend, but there was a significant decline in the megalopa stage (Fig. S10 D, E, F).

**Fig. 6.**
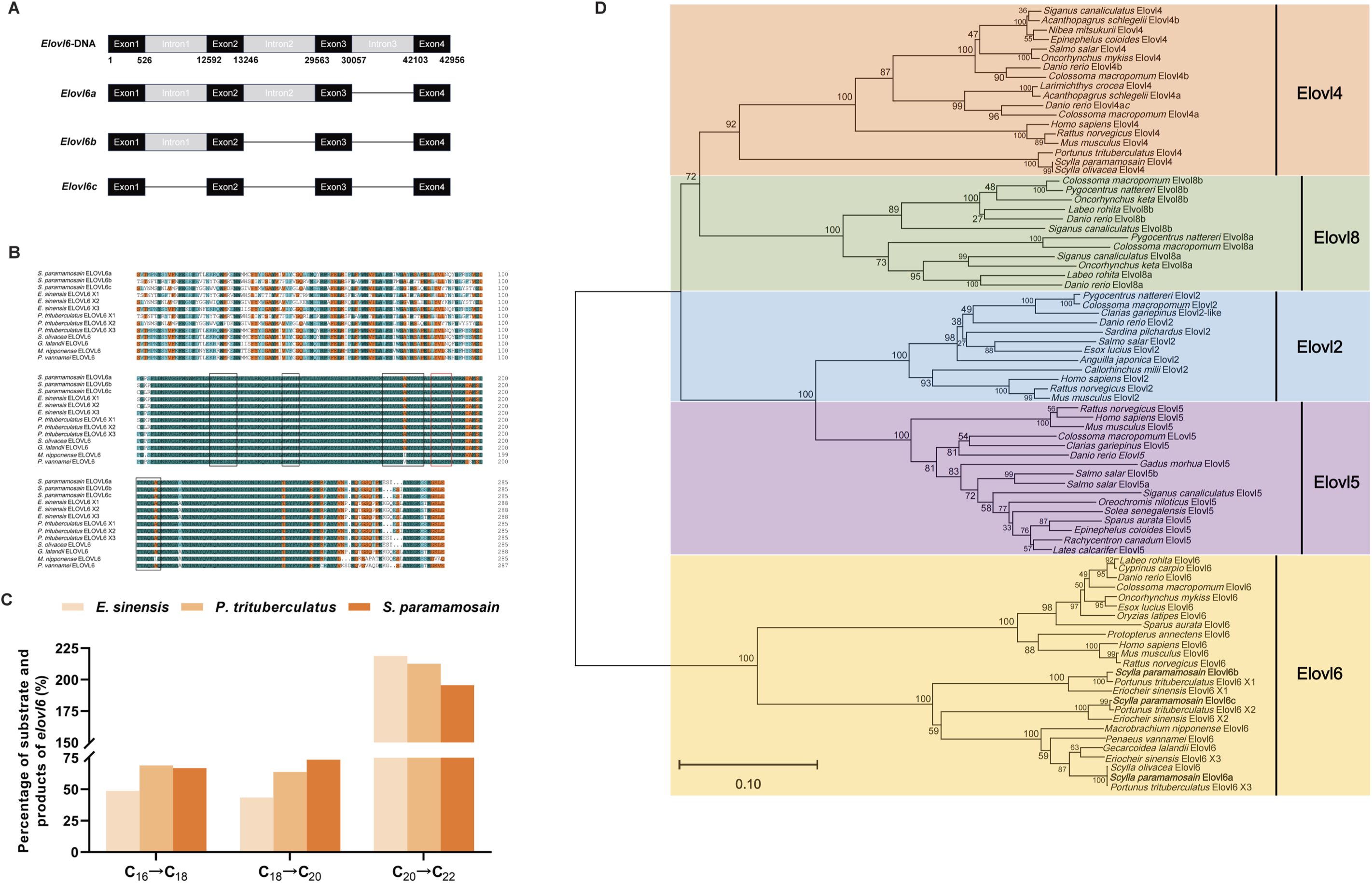
Neo-functionalization of the *elovl6* gene in the LC-PUFA synthesis pathway in crustacean species. **(A)** Schematic diagram of the alternative splicing in *elovl6*. *elovl6*-DNA, the DNA sequence of *elovl6*; *elovl6a*, the *elovl6* transcript containing the first and second introns; *elovl6b*, the *elovl6* transcript containing the first intron; *elovl6c*, the *elovl6* transcript of introns normal splicing. Black boxes represent exons. Gray boxes represent introns. **(B)** Alignment of the amino acid sequences of *elovl6* from crustacean species, including *Scylla olivacea* (QEV88898.1), *Gecarcoidea lalandii* (QKG32709.1), *Macrobrachium nipponense* (ANM86278.1), and *Penaeus vannamei* (AKJ77890.1). Identical amino acid residues were indicated by dark green, and similar residues were indicated by blue and orange. The conserved HXXHH histidine motif was in the box, and highly conserved motifs (KXXEXXDT, NXXXHXXMYSYY, KXXKXX, and TXXQXXQ) were framed in a black box. **(C)** Phylogenetic tree comparing *S. paramamosain* Elovl6 with elongase proteins from other organisms. The tree was constructed using the neighbour-Joining method by MEGA 11.0.13. The horizontal branch length is proportional to amino acid substitution rate per site. The numbers represent the frequencies with which the tree topology presented was replicated after 1000 iterations. **(D)** Comparison of Elovl6 extended carbon chain conversion efficiency in *E. sinensis*, *P. trituberculatus*, and *S. paramamosain*.

Furthermore, *S. paramamosain* Elovl6a, Elovl6b, and Elovl6c were functionally characterized through a yeast heterologous expression system (Table S5). Upon addition of 16:0 fatty acid substrate, the Elovl6-transformed yeast exhibited the production of elongation products consisting of saturated fatty acids (SFA) ranging from C_18_ to C_22_. Similarly, incubation with 18:1n-9 resulted in the generation of several elongation products of monounsaturated fatty acids (MUFA). Endogenous 18:1n-9 was also elongated to form 20:1n-9 and subsequently further elongated to produce 22:1n-9. Regarding PUFA substrates, all C_18_ PUFA substrates tested were significantly converted to their respective C_20_ products, with elongation observed for 18:3n-3, 18:4n-3, 18:2n-6, and 18:3n-6 to their corresponding C_20_ PUFA. Notably, *S. paramamosain* Elovl6 exhibited a higher affinity toward n-3 PUFA than n-6 PUFA when utilizing C_18_ PUFA substrates. However, neither of the C_20_ LC-PUFA, arachidonic acid (ARA; 20:4n-6) or eicosapentaenoic acid (EPA; 20:5n-3), underwent elongation and so 22:4n-6 and 22:5n-3, respectively, were not generated. Additionally, a significant decrease in the expression level of the *elovl6* gene was observed following 18 h and 36 h interference in isolated hepatopancreas of green mud crab (Fig. S10G), resulting in a notable reduction in 16:0, 16:1n-7, 18:1n-9, 18:2n-6, 18:3n-3, 20:5n-3, and 22:6n-3. This further validated the capability of Elovl6 for elongation of C_18_ PUFA to C_20_ PUFA (Fig. S10H). Thus, the capacity of Elovl6 of *S. paramamosain* for elongation of C_18_ PUFA, in addition to SFA and MUFA, was confirmed. In contrast, Elovl6 proteins in mammals (Matsuzaka et al, 2007; Junjvlieke et al, 2019) and teleosts (Li et al, 2020; Wang et al, 2020) are only involved in the elongation of SFA and MUFA as they have only been demonstrated to elongate 16:0 and 16:1 to 18:0 and 18:1, respectively, with other potential activities of *elovl6* gene products not yet reported.

The aforementioned results indicate that Elovl6 may undergo neofunctionalization in the Decapoda species *S. paramamosain*. Decapod crabs inhabit a diverse range of environments, including seawater (e.g., *P. trituberculatus*), freshwater (e.g., *E. sinensis*), and brackish water (e.g., *S. paramamosain*). It was reported that the multifunctionalization of existing genes and neo-functionalization of duplicated genes have occurred in catadromous and freshwater teleost species such as flatfish family Achiridae (Matsushita et al, 2020) and freshwater three-spined stickleback (Ishikawa et al, 2021). To investigate the influence of the environment on the evolutionary adaptation of key enzymes of LC-PUFA synthesis in Decapoda crab species, three transcripts of the *E. sinensis and P. trituberculatus elovl6* genes were functionally characterized. The findings demonstrated that the *Ptelovl6* and *Eselovl6* genes exhibited comparable fatty acid elongation activities to that of *Spelovl6* (Fig. 6C), implying the potential neofunctionalization of Elovl6 in decapod crabs. The phylogenetic analysis of *elovl6* in *E. sinensis*, *P. trituberculatus*, and *S. paramamosain* showed the evolutionary difference between *elovl6* and *elovl2/*4/5/8 (Fig. 6D), which indicated that neo-functionalization of the *elovl6* gene in the LC-PUFA synthesis pathway in crustaceans occurred.

## Discussion

Crabs represent a pivotal model system for exploring the molecular underpinnings of adaptive evolution within Decapoda. Here, we present a comprehensive analysis of the green mud crab *S. paramamosain* genome using Nanopore ultralong-read sequencing, achieving remarkable contig N50 and scaffold N50 values that substantially surpass previous assemblies (Zhang et al, 2019; Song et al, 2016; Tang et al, 2021; Tang et al, 2020; Zhao et al, 2021; Cui et al, 2021; Chebbi et al, 2019). By anchoring contigs to 49 pseudochromosomes, we attained a more precise depiction of the genome architecture, with Chr06 revealing a prominent sex-linked signal, offering foundational insights into sex determination mechanisms in decapod crustaceans. Furthermore, the detection of potential ancient whole-genome duplication events suggests a dynamic evolutionary history underlying the genomic organization of *S. paramamosain*.

The highly repetitive nature of crustacean genomes presents significant challenges in maintaining genome integrity (Zhang et al, 2019; Polinski et al, 2021). Our investigation into genome assembly emphasized the impact of ultralong-read sequencing in resolving repetitive sequences, culminating in the identification of 58.97% of the genome as repetitive. Repetitive sequences and TEs significantly contribute to genome expansion and evolution in *S. paramamosain*, with a higher proportion observed compared to several other crustacean species (Zhang et al, 2019; Song et al, 2016; Tang et al, 2021; Tang et al, 2020; Cui et al, 2021; Chebbi et al, 2019). The identification of numerous SSRs and TRs underscores the importance of these elements in shaping the genomic landscape of *S. paramamosain*. Analysis of SSRs, TRs, and TEs provided insights into genome expansion and evolution. *S. paramamosain* exhibits 33,662 protein-coding genes, highlighting gene richness compared to other crustaceans (Zhang et al, 2019; Song et al, 2016; Tang et al, 2021; Tang et al, 2020; Zhao et al, 2021; Cui et al, 2021; Chebbi et al, 2019). The annotation of these genes to various databases enhances our understanding of green mud crab genomics. Gene family analysis revealed species-specific genes enriched in pathways related to development, indicating potential adaptations. Phylogenetic analysis positioned green mud crabs among crustaceans, highlighting their evolutionary divergence.

The investigation of Hox genes in *S. paramamosain* elucidated their role in directing tissue differentiation and morphological development during larval stages, particularly in the development of pleopods. The expression patterns of Hox genes during different larval stages suggest their involvement in the regulation of key developmental processes, with potential implications for understanding the evolution of crustacean body plans. We investigated the regulatory mechanisms of Hox genes in pleopod development, demonstrating the crucial role of *Abd-A* in abdominal limb development. Posttranscriptional regulation involving novel-miR1317 fine-tunes *Abd-A* levels, unveiling a nuanced layer of gene regulation.

Additionally, our exploration of the *fru* gene in ovarian development underscored its potential role in female gonadal development and maturation. The identification of novel-miR35 as a potential regulator of *Spfru2* adds complexity to the gene regulatory network governing reproductive processes. The divergent expression patterns of *fru* in male and female crabs, alongside its localization in gonadal tissues, highlight its potential involvement in ovarian maturation and gonadal development. Furthermore, the identification of novel microRNA targeting *fru* underscores the intricate posttranscriptional regulatory mechanisms governing reproductive processes in crustaceans.

The neofunctionalization of the elovl6 gene in the long-chain polyunsaturated fatty acid (LC-PUFA) synthesis pathway represents a key adaptation in *S. paramamosain*. Here, we elucidate the neo-functionalization of the *elovl6* gene in the synthesis of LC-PUFA, shedding light on its multiple transcripts and tissue-specific expression. Comparative analysis with *S. paramamosain*, *P. trituberculatus*, and *E. sinensis* suggests evolutionary divergence in LC-PUFA biosynthesis pathways, potentially driven by environmental adaptation.

Overall, our findings shed light on various aspects of crab biology, including genome sequencing, assembly, and annotation, as well as gene family expansion, contraction, and regulatory mechanisms governing crucial developmental processes such as metamorphosis, reproductive strategies, and fatty acid metabolism. The present findings significantly advance our understanding of the green mud crab genome, providing a valuable resource for future studies in comparative genomics, evolutionary biology, and functional genomics in crustaceans. Future studies leveraging this genomic resource will further elucidate the genetic basis of key adaptive traits and inform conservation and management strategies for economically important crab species.

## Methods and materials

### Sample preparation and genomic DNA isolation

Genomic DNA for sequencing was extracted from the testis of a wild adult male green mud crab *S. paramamosain* with stage III testis, which was caught off the coast of Shantou, China. A Grandomics Genomic BAC-long DNA Kit was used to isolate ONT ultralong DNA according to the manufacturer’s guidelines. The total DNA quantity and quality were evaluated using a NanoDrop One UV-Vis Spectrophotometer (Thermo Fisher Scientific, Waltham, MA) and a Qubit 3.0 Fluorometer (Invitrogen Life Technologies, Carlsbad, CA). Large DNA fragments were obtained through gel cutting with the Blue Pippin system (Sage Science, Beverly, MA). Qualification of DNA involved visual inspection for foreign matter, assessment of degradation and size *via* 0.75 % agarose gel electrophoresis, check on purity (OD260/280 between 1.8-2.0; OD260/230 between 2.0-2.2) using Nanodrop, and precise quantification with Qubit 3.0 Fluorometer (Invitrogen, USA).

### Library construction and sequencing

Approximately 8-10 µg of genomic DNA (> 50 kb) was selected using the SageHLS HMW library system (Sage Science, Beverley MA, USA) and processed with the Ligation Sequencing 1D Kit (Oxford Nanopore Technologies, Shanghai, China) following the manufacturer’s instructions. Library construction and sequencing were conducted on the PromethION (Oxford Nanopore Technologies) at the Genome Center of Grandomics (Wuhan, China). After quality inspection, large DNA fragments were recovered using the BluePippin automatic nucleic acid recovery instrument. Terminal repair, A-tailing, and ligation were performed using the LSK109 connection kit. Qubit was employed to assess the constructed DNA library precisely. The library was loaded into a flow cell, transferred to the PromethION sequencer, and subjected to real-time single-molecule sequencing. In the ONT sequencing platform, base calling, the conversion of nanopore-generated signals to base sequences (Wick et al, 2019), was executed using the Guppy toolkit (Oxford Nanopore Technologies). Pass reads with a mean qscore_template value greater than or equal to 7 were obtained and directly used for subsequent assembly (https://github.com/nanoporetech/taiyaki).

### Genome assembly, evaluation and correction

Filtered reads post quality control were employed for a pure three-generation assembly using NextDenovo software (reads_cutoff:1k, seed_cutoff:28k) (https://github.com/Nextomics/NextDenovo.git). The NextCorrect module was employed to correct the original data, yielding a consistency sequence (CNS sequence) after 13 Gb of error correction. *De novo* assembly using the NextGraph module produced a preliminary assembly of the genome. ONT three-generation data and Pb HiFi three-generation data were utilized with Nextpolish software (https://github.com/Nextomics/NextPolish.git) for genome correction. The corrected genome (Polish Genome) was obtained after three rounds of correction for both ONT and Pb HiFi data. Bwa mem default parameters were used to compare next-generation data to the genome, and Pilon was iteratively calibrated three times to derive the final genomic sequence. GC depth analysis and BUSCO prediction (https://busco.ezlab.org/) were used to assess genome quality and completeness.

### Chromosome anchoring by Hi-C sequencing

To anchor hybrid scaffolds onto chromosomes, genomic DNA for Hi-C library construction was extracted from green mud crab testes. Cell samples from the tissue used for genomic DNA sequencing were employed for Hi-C library preparation. The process involved cross-linking cells with formaldehyde, lysing cells, resuspending nuclei, and subsequent steps leading to proximity ligation. After overnight ligation, cross-linking was reversed, and chromatin DNA manipulations were performed. DNA purification and shearing to 400 bp lengths were followed by Hi-C library preparation using the NEBNext Ultra II DNA library Prep Kit for Illumina. Sequencing on the Illumina NovaSeq/MGI-2000 platform completed the Hi-C procedure. Briefly, cell samples were fixed with formaldehyde and subjected to lysis and extraction for sample quality assessment. After passing the quality test, the Hi-C fragment preparation process involved chromatin digestion, biotin labeling, end ligation, DNA purification, and library construction. The library was sequenced on the MGI-2000 platform, and data were processed to extract high-quality reads. The analysis included filtering for adapters, removing low-quality reads, and eliminating reads with an N content exceeding 5. Reads were aligned using Bowtie2 (Langmead and Salzberg 2012), and contig clustering was performed using LACHESIS software (Joshua and Burton 2013).

### Gene annotation

Tandem repeats were annotated using GMATA (https://sourceforge.net/projects/gmata/?source=navbar) and Tandem Repeats Finder (TRF) (http://tandem.bu.edu/trf/trf.html), identifying simple repeat sequences (SSRs) and all tandem repeat elements. Transposable elements (TEs) were identified through an *ab initio* and homology-based approach, with RepeatMasker (https://github.com/rmhubley/RepeatMasker) used for searching known and novel TEs. Gene prediction employed three methods: GeMoMa (http://www.jstacs.de/index.php/GeMoMa) for homolog prediction, PASA (https://github.com/PASApipeline/PASApipeline) for RNAseq-based prediction, and Augustus (https://github.com/Gaius-Augustus/Augustus) for *de novo* prediction. EVidenceModeler (EVM) (http://evidencemodeler.github.io/) integrated gene sets, which underwent further filtering for transposons and erroneous genes. UTRs and alternative splicing regions were determined using PASA based on RNA-seq assemblies. Functional annotation involved comparisons with public databases, including SwissProt, NR, KEGG, KOG, and Gene Ontology. InterProScan identified putative domains and GO terms. BLASTp (https://blast.ncbi.nlm.nih.gov/Blast.cgi) against public protein databases was used to assess gene function information. The noncoding RNA (ncRNA) prediction entailed using tRNAscan-SE (http://lowelab.ucsc.edu/tRNAscan-SE/) and Infernal cmscan (http://eddylab.org/infernal/) for tRNAs and other noncoding RNAs. BUSCO was employed for gene prediction evaluation, aligning annotated protein sequences to evolution-specific BUSCO databases.

### Evolutionary analysis

The evolutionary analysis entailed the examination of gene families, construction of phylogenetic trees, estimation of divergence times, exploration of gene expansion and contraction phenomena, identification of orthologous genes, scrutiny of positively selected genes, and investigation of whole-genome duplications. To ascertain homologous relationships between *S. paramamosain* and other animal species, protein sequences were acquired and aligned using OrthMCL (https://orthomcl.org/orthomcl/). Initially, protein sets were gathered from 14 sequenced animal species, and the longest transcripts for each gene were selected, excluding miscoded and prematurely terminated genes. Subsequently, pairwise alignment of these extracted protein sequences was conducted to identify conserved orthologs, employing Blastp with an E-value threshold of ≤ 1 × 10^-5^. Further identification of orthologous intergenome gene pairs, paralogous intragenome gene pairs, and single-copy gene pairs was achieved using OrthMCL. Species-specific genes, encompassing *S. paramamosain*-specific unique genes and unclustered genes, were extracted. Functional annotation and enrichment tests of species-specific genes were performed utilizing information from homologs in the online Gene Ontology (http://www.geneontology.org/) and KEGG (Kyoto Encyclopedia of Genes and Genomes) (https://www.genome.jp/kegg/) databases.

Building upon the orthologous gene sets identified with OrthMCL, molecular phylogenetic analysis was executed using shared single-copy genes. Coding sequences were extracted from single-copy families, followed by multiple alignment of each ortholog group using MAFFT (https://mafft.cbrc.jp/alignment/software/). Gblocks were applied to eliminate poorly aligned sequences, and the GTRGAMMA substitution model of RAxML (https://cme.h-its.org/exelixis/web/software/raxml/hands_on.html) was employed for phylogenetic tree construction with 1000 bootstrap replicates. The resulting tree file was visualized using Figtree/MEGA (http://tree.bio.ed.ac.uk/software/figtree/). The RelTime tool (https://www.megasoftware.net/) of MEGA-CC was then utilized to compute mean substitution rates along each branch and estimate species divergence times, with three fossil calibration times obtained from the TimeTree (http://www.timetree.org/) database serving as temporal controls, including the divergence times of *S. paramamosain*.

The detection of significant expansions or contractions in specific gene families, often indicative of adaptive divergence in closely related species, was carried out based on OrthoMCL results. CAFE (https://github.com/hahnlab/CAFE), employing a birth and death process to model gene gain and loss over a phylogeny, was used for this purpose. Furthermore, in accordance with the neutral theory of molecular evolution, the ratio of the nonsynonymous substitution rate (Ka) to the synonymous substitution rate (Ks) of protein-coding genes was calculated. The average Ka/Ks values were determined, and a branch-site likelihood ratio test using Codeml (http://abacus.gene.ucl.ac.uk/software/) from the PAML package was conducted to identify positively selected genes within the *S. paramamosain* lineage. Genes with a p value < 0.05 under the branch-site model were considered positively selected.

Whole-genome duplication events in the green mud crab genome were investigated using four-fold synonymous third-codon transversion (4DTv) and synonymous substitution rate (Ks) estimation. Initial steps involved extracting protein sequences and conducting all-vs.-all paralog analysis through self-Blastp in these plants. After filtering by identity and coverage, the Blastp results underwent analysis with MCScanX (Wang et al, 2012), and the respective collinear blocks were identified. Subsequently, Ks and 4DTV were calculated for the syntenic block gene pairs using KaKs-Calculator (https://sourceforge.net/projects/kakscalculator2/), and potential WGD events in each genome were evaluated based on their Ks and 4DTv distribution.

### RNA-seq sample collection and sequencing

A total of 1 μg of high-quality RNA per experimental group (including different developmental stages, different tissues, and gonads from different mating periods of *S. paramamosain*) was allocated for RNA sample preparations. Sequencing libraries were constructed using the NEBNext UltraTM RNA Library Prep Kit for Illumina (NEB, USA), following the manufacturer’s guidelines. Subsequently, each sample was labeled with index codes for sequence characterization. In brief, mRNA was isolated from total RNA using poly T oligomer magnetic beads. NEBNext First-Strand Synthesis Reaction Buffer (5X) was employed, utilizing bivalent cations for high-temperature cleavage. First-strand cDNA was synthesized with random hexamer primers and M-MuLV Reverse Transcriptase. Subsequently, the second strand of cDNA was synthesized using DNA Polymerase I and RNase H, and any remaining overhangs were converted to blunt ends by exonuclease/polymerase. Following adenylation of the 3’ end of DNA fragments, the NEBNext adaptor with a hairpin loop structure was ligated for hybridization. Library fragments, preferentially selecting 240 bp cDNA fragments, were purified using the AMPure XP system (Beckman Coulter, Beverly, USA). The USER enzyme (NEB, USA) was introduced to the cDNA of the selected size and adaptor, and PCR was conducted with Phusion High-fidelity DNA polymerase after 15 min at 37 ℃ and 5 min at 95 ℃. Universal PCR primers and index (X) primers were utilized for PCR. Finally, the PCR products were purified using the AMPure XP system, and the library quality was assessed using the Agilent Bioanalyzer 2100 system. Following the manufacturer’s recommendations, the TruSeq PE Cluster Kit v4-cBot -HS (Illumina) was employed to cluster the index-coded samples on the cBot Cluster Generation System. After clustering, library preparations were sequenced on the Illumina platform to generate paired-end reads.

### Microinjection and RNA interference

Primer design for the amplification of the *Abd-A*, *fru2* and *elovl6* gene fragments and subsequent dsRNA synthesis was based on the gene sequences. Specific primers were designed incorporating protective bases and T7 promoter sequences at the 5’ end of both the positive and negative primers to facilitate dsRNA synthesis. The plasmid containing the *Abd-A* and *elovl6* gene sequence fragments served as the template for PCR amplification, and the resulting product underwent purification. The PCR product, now equipped with the T7 promoter, was utilized as the template for the synthesis of dsRNA, a crucial step for gene interference through *in vitro* transcription. Subsequently, zoeal stage IV of the green mud crab were positioned on a custom-made agarose gel plate. Injection of dsRNA and the injection indicator mixture was carried out at the cuticle and abdominal space of the larvae using a microinjector. As the larvae progressed to the zoea V stage, the development of the abdomen and limbs in the juveniles was meticulously observed through electron microscopy.

### Hepatopancreas treatment *in vitro*

Crabs were anesthetized on ice for 10 min, followed by sterilization in 75 % ethanol for 5 min. Hepatopancreas tissues were subsequently dissected and first infiltrated with phosphate buffered saline (PBS) containing 1 % penicillin and streptomycin for 30 min, then washed 8 times for 5 min each time using PBS solution as above. Next, hepatopancreas tissues were chopped by scissors into fragments of approximately 20 mg. The fragments were then precultured at room temperature (25 ℃) in a 24-well-plate with 150 µL of Leibovitz L15 medium (containing 1% penicillin and streptomycin). The 24-well culture plates were supplemented with dsRNA-Elovl6 and dsRNA-EGFP (control group) and placed in a 28 ℃ incubator for cultivation. After 18 h, 24 h, and 36 h of culture, tissue fragments were collected from each treatment for total RNA extraction and subsequent qRT-PCR analysis, with corresponding parallel samples of hepatopancreas tissue also collected for fatty acid analysis.

### Sequence and phylogenetic analysis

To provide important clues for predicting functions or evolution of the *elovl6* genes, multiple sequence alignments were performed with the DNAMAN software (v6.0.3.99, Lynnon Corporation, USA). The phylogenetic tree mapped using the neighbor-Joining method with MEGA software (v11.0.13, Arizona State University, USA), on the basis of deduced amino acid sequences of Elovls from *S. paramamosain* and representative mammals, teleosts, and all functionally characterized or uncharacterized Elovl from crustacean species. Confidence in the resulting phylogenetic tree branch topology was measured by bootstrapping through 1000 replications.

### Quantitative real time PCR (qRT-PCR)

Using the miRcute miRNA Isolation Kit, miRNAs were isolated from distinct developmental stages and tissues of *S. paramamosain*. Subsequently, the commercially accessible miRcute Enhanced miRNA cDNA First Strand Synthesis Kit was deployed for the reverse transcription process. Finally, the miRcute Enhanced miRNA Fluorescence Quantitative Detection Kit was applied in the qPCR analysis to elucidate the expression dynamics of both miRNAs and the genes. The total RNA extraction was extracted using the RNAiso Plus kit (Takara Co. Ltd., Japan). Preceding the quantitative real-time polymerase chain reaction (qRT-PCR), the RNA samples underwent treatment with RQ1 RNase-Free DNase (Takara Co. Ltd.) to eradicate genomic DNA contamination. Subsequently, cDNA synthesis was conducted using 500 ng of DNase-treated RNA and the Talent qPCR Premix (SYBR Green) kit (TIANGEN Biotech Co., Ltd., Beijing), in accordance with the manufacturer’s introductions. The qPCR primers were meticulously designed using Primer 6.0 software (Table S6). The qRT-PCR reactions were performed employing a Mini Option real-time detector (Roche LightCycle@480). Each reaction mixture comprised 10 µL of Talent qPCR Premix (2×), 0.6 µL of PCR forward primer (10 µM), 0.6 µL of PCR reverse primer (10 µM), 2.0 µL of RT reaction solution containing cDNA at an amount of 20 ng, and 6.8 µL of RNase-free water. The amplification protocol involved an initial denaturation step at 95 °C for 3 min, followed by 40 cycles of denaturation at 95 °C for 5 s, annealing at 60 °C for 10 s, and extension at 72 °C for 15 s. A subsequent melting curve analysis ensued by sustaining the reaction at 72 °C for 6 s, followed by a concise denaturation step at 95 °C for 5 s. For verification of correct amplicon sizes, all products were initially resolved through agarose gel electrophoresis. Normalization of gene expression levels to the reference gene (*18s rRNA*) was conducted. The gene expression levels were calculated using the optimized comparative Ct method (2^-ΔΔCt^).

### Protein expression detection using Western blotting

Protein extraction from cephalothorax and abdomen samples of green mud crab larvae was performed using a cryogenic protein lysis solution following flash freezing with liquid nitrogen. Subsequently, total protein was extracted from each sample. After quantifying the protein concentration using the BCA protein assay kit (Sangon Biotech Co. Ltd., Shanghai, China), an equal number of proteins were loaded for SDS-PAGE electrophoresis. Following separation based on molecular weight, the proteins were transferred onto a PVDF membrane from the polyacrylamide gel and subsequently probed with specific antibodies. Subsequent steps included incubation with HRP-conjugated secondary antibodies and detection using an ECL hypersensitive chemiluminescence kit (Sangon Biotech).

### Fluorescence *in situ* hybridization (FISH)

The FISH analysis was performed on *S. paramamosain* gonads, which were fixed in 4 % paraformaldehyde prepared with DEPC-treated water two hours prior to sectioning at a thickness of 10-12 µm using a cryostat set at -20 to -25 °C. Subsequently, no more than three sections per slide were thaw-mounted onto charged Superfrost Plus slides. The *fru2* probe, modified with FITC fluorescence at the 3′ end, was synthesized commercially (Sangon Biotech). The prehybridization and hybridization procedures followed the method described by Pinaud et al (2008). Initially, a prehybridization solution diluted with high-grade formamide in a 1:1 ratio was added to each slide and coverslip section (50 µL per sample). Then, the slides were placed in a humid chamber for 30 min at room temperature before removing the coverslips in a 2× SSC solution. After prehybridization, the probe was added to the hybridization buffer at a concentration of 1 ng/µL. Twenty-five mL hybridization mix was applied to each slide and coverslip section before placing them in a metal slide holder immersed in a mineral oil bath maintained at 65 °C for 14-16 h. Following hybridization, the slide holder was removed from the mineral oil bath and slides were rinsed twice (30 s each) in chloroform followed by immersion in a fresh batch of 2× SSC solution without coverslips for washing at room temperature for 1 h; subsequently washed again in an SSC solution containing 50 % formamide at 65 °C for 1 h; finally washed twice (each time lasting 30 min) in ten-fold diluted (0.1×) SSC solution while maintaining the temperature at 65 °C throughout these washes. The signals resulting from hybridization on each slide were ultimately detected using fluorescence microscopy.

### Yeast expression and fatty acid detection

PCR amplification of open reading frames (ORFs) corresponding to *Eselovl6*, *Ptelovl6*, and *Spelovl6* was performed using a high-fidelity DNA polymerase, KOD-Plus-Neo (Toyobo, Japan), and cDNA templates. Specific primers that incorporated BamHI (GGATCC) and EcoRI (GAATTC) restriction sites were employed in accordance with the manufacturer’s instructions. The primers designed for ORF cloning, with their respective restriction sites, are detailed in Table S6. For *Eselovl6* and *Ptelovl6*, the amplification process involved an initial denaturation step at 96 °C for 3 min, followed by 23 cycles of denaturation at 95 °C for 30 s, annealing at 58 °C for 15 s, and extension at 72 °C for 20 s, culminating in a final extension at 72 °C for 1 min. The PCR conditions for *Spelovl6* included an initial denaturing step at 94 °C for 2 min, followed by 35 cycles of denaturation at 98 °C for 30 s, annealing at 94 °C for 30 s, extension at 68 °C for 30 s, and a final extension at 72 °C for 7 min.

Subsequently, the resulting DNA fragments were purified using the TIAN quick mini purification kit (Tiangen Biotech). After purification, the fragments were digested with the corresponding BamHI and EcoRI restriction endonucleases (Thermo Scientific, USA) and ligated into a similarly restricted pYES2 yeast expression vector (Invitrogen, USA) using the DNA Ligation Kit Mighty Mix (Takara, Japan). The resulting plasmid constructs, namely, *pYEselovl6*, *pYPtelovl6*, and *pYSpelovl6*, were then introduced into *E. coli* DH5α competent cells (Takara, Japan), which were subsequently screened for the presence of recombinants *via* PCR.

The transformation and selection of yeast with recombinant plasmids, as well as yeast culture, followed modified protocols based on a previous study (Monroig et al, 2012). In summary, purified plasmids, inserted with either one of the elovl6 ORFs or no insert, were used to transform yeast (*Saccharomyces cerevisiae* strain INVSc1) competent cells through the polyethylene glycol/lithium acetate (PEG/LiAc) method. The transformed yeast containing the corresponding recombinants were selected on *S. cerevisiae* minimal solid media plates without uracil (SC-Ura, containing 2 % glucose and 2 % yeast agar) and cultured at 30 °C for 3 d. Single colonies were then picked and transferred into liquid SC-Ura medium with 2 % glucose, followed by further growth in a shaking incubator at 30 °C under 250 rpm for 2 d. Yeast genomic DNA was extracted from the bacterial solution using a yeast genomic DNA rapid extraction kit (Solarbio, Beijing, China), and the presence of the resulting plasmid constructs in *S. cerevisiae* was identified and confirmed by PCR screening.

For heterologous expression, successfully transformed yeast was initially cultured for 24 h in SC-Ura broth containing 2 % raffinose at 30 °C and 250 rpm. Cultured *S. cerevisiae* cells were then diluted to an OD600 of 0.4 in SC-Ura broth and allowed to grow until the OD600 reached 1. The suspensions were centrifuged to obtain precipitated yeast, which was then resuspended in SC-Ura broth (containing 2 % galactose) at an OD600 of 0.4. Next, a mixture of the suspensions, 10 % tergitol-type (NP-40, Beyotime Biotechnology, Shanghai, China), and corresponding fatty acid substrates substrates (including 16:0, 18:1n-9, 18:2n-6, 18:3n-3, 18:3n-6, 18:4n-3, 20:4n-6, and 20:5n-3) at final concentrations of 0.50 mM for C_16_, 0.50 mM for C_18_, and 0.75 mM for C_20_ were added to the centrifuge tubes. The tubes were placed into a shaker and incubated at 30 °C for 2 d at 250 rpm. Subsequently, the yeast was collected *via* centrifugation, washed with Hanks’s balanced salt solution (Solarbio), and processed for fatty acid analysis. This experiment was replicated using three separate recombinant colonies for each recombinant yeast strain.

### Fatty acid analysis by GC-MS

The yeast obtained above underwent a 48-h freeze-drying process and was subsequently ground into powder. Approximately 200 mg of yeast powder was then thawed at 4 °C and placed in a 12-mL screw-top glass tube with a Teflon-sealed lid and 0.25 mg/mL methylation solution, which was composed of 99 mL methanol, 1 mL sulfuric acid, and 0.025 g butylated hydroxy toluene as antioxidant, was added to the tube. After vortexing for 2 min and ultrasonic disruption for 30 min, the samples were incubated in a water bath at 80 °C for 4 h. After cooling, 1 mL of n-hexane was added to the mixture and shaken vigorously for 1 min, and 1 mL of ultrapure water then added to facilitate layer separation. The resulting supernatant was then filtered through a nylon syringe filter with a 0.22 µm ultrafiltration membrane (SCAA-104, ANPEL, China) and collected into a clean ampoule. The fatty acid methyl ester (FAME) solution was concentrated under a stream of nitrogen gas in a Termovap sample concentrator and the FAME resuspended in 600 µL of n-hexane and stored at -20 °C until analysis. The FAME samples were separated and analyzed by gas chromatography‒mass spectrometry (GC‒ MS) using an Agilent 7890B-5977A GC-MS (Agilent Technologies, Santa Clara, CA, USA) equipped with a fused-silica ultra inert capillary column (DB-WAX, 30 m × 250 µm internal diameter, film thickness 0.25 µm; Agilent J & W Scientific, CA, USA). The oven temperature was initially increased from 100 ℃ to 200 ℃ at a rate of 10 ℃/min, with a hold time of 5 min at 200 ℃. Subsequently, the temperature was increased to 230 ℃ at 2 ℃/min with a hold time of 10 min at 230 ℃, followed by a final ramp from 230 to 240 at 10 ℃/min. The injection, interface, and ion source temperatures were adjusted to 250, 240, and 230 °C, respectively. High-purity helium (99.999 %) served as the carrier gas with a constant flow rate of 1 mL/min. A 0.5 µL sample was injected at a 1:20 split ratio by an autosampler. The collision energy was set at 70 eV, and mass spectra data were acquired in full scan mode (scanning range 40–500 m/z). Fatty acids were identified through mass spectrometry, referencing a commercially available standard library (National Institute of Standards and Technology Mass Spectral Library 2011) and the relative retention times of standards. The elongation conversion efficiencies of the respective added fatty acid substrates were determined by calculating the proportion of exogenously added fatty acids (FAs) to the elongated FA products, given by the following equation.

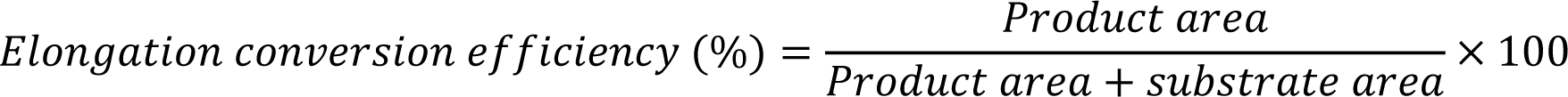

## Data availability

The whole genome sequences are deposited at DDBJ/ENA/GenBank under the accession JAYKKS000000000. The version described in this paper is version JAYKKS010000000. Raw sequencing data are deposited in NCBI under BioProject PRJNA1059155.

## Acknowledgements

This study was supported by the National Key Research & Development Program of China (2018YFD0900201), the National Natural Science Foundation of China (42076133, 42306126, 42306125), the National Plan for the Special Support for Top-notch Talents (ZUTINGZI201548), the Leading Talent Project of Special Support Plan of Guangdong Province (2019TX05N067), the Guangdong Natural Science Foundation (2022A1515110488, 2022A1515111151), and the STU Scientific Research Foundation for Talents (NTF21016, NTF21020). We express our gratitude to Dr. Noah Esmaeili for his valuable input in enhancing the linguistic quality of the manuscript.

## Author contributions

Hongyu Ma provided the funding of the research and conceptualized the whole experiment. Yin Zhang, Ye Yuan, Mengqian Zhang, Xiaoyan Yu, Bixun Qiu and Fangchun Wu collected the specimens. Yin Zhang, Ye Yuan, Xiaoyan Yu, Bixun Qiu, Fangchun Wu, Jiajia Zhang and Mengqian Zhang performed the experiment. Yin Zhang, Ye Yuan and Mengqian Zhang analyzed the data and prepared the required figures. Yin Zhang, Ye Yuan and Mengqian Zhang wrote the manuscript. Shaopan Ye and Yin Zhang upload the genome data. Hongyu Ma, Shaopan Ye, Wenxiao Cui, Jonathan Y. S. Leung, Mhd Ikhwanuddin, Tariq Dildar, Waqas Waqas and Douglas R. Tocher revised the manuscript and contributed to additional discussion. All authors read and approved the final manuscript.

## Additional information

Supplementary information is available for this paper.

Correspondence and requests for materials should be addressed to Hongyu Ma.

