## Supplementary material for "High-resolution chromosome-level genome provides molecular insights into adaptive evolution in crabs": Suplementary tables and figures

### Supplementary Figures

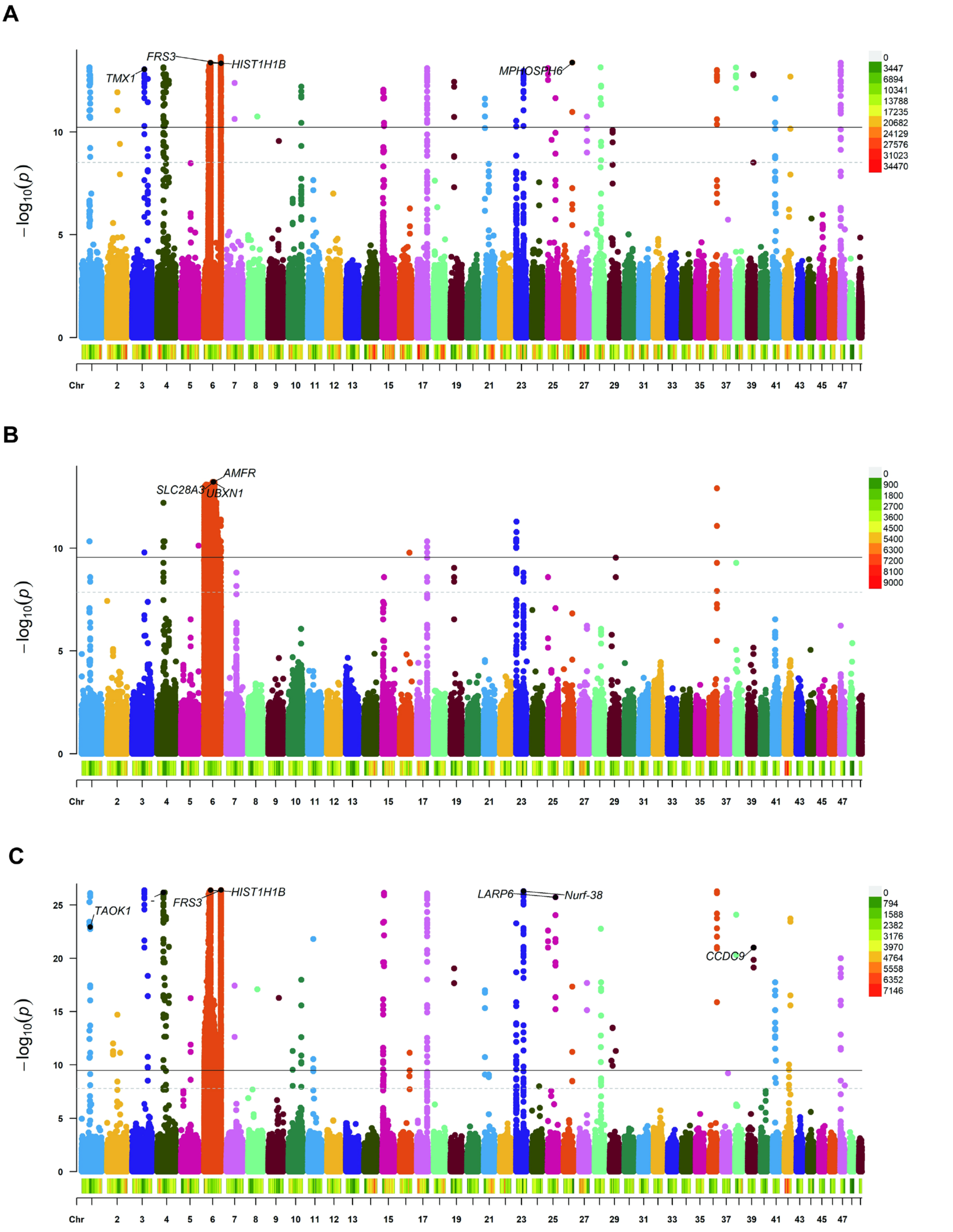


**Figure S1** Genome-wide association studies (GWAS) analysis of *Scylla paramamosain* genome using different generations of populations. **(A)** 146 individuals from wild. **(B)** 130 individuals from second generation of family group. **(C)** Complex of 146 individuals from wild and 130 individuals from second generation of family group.

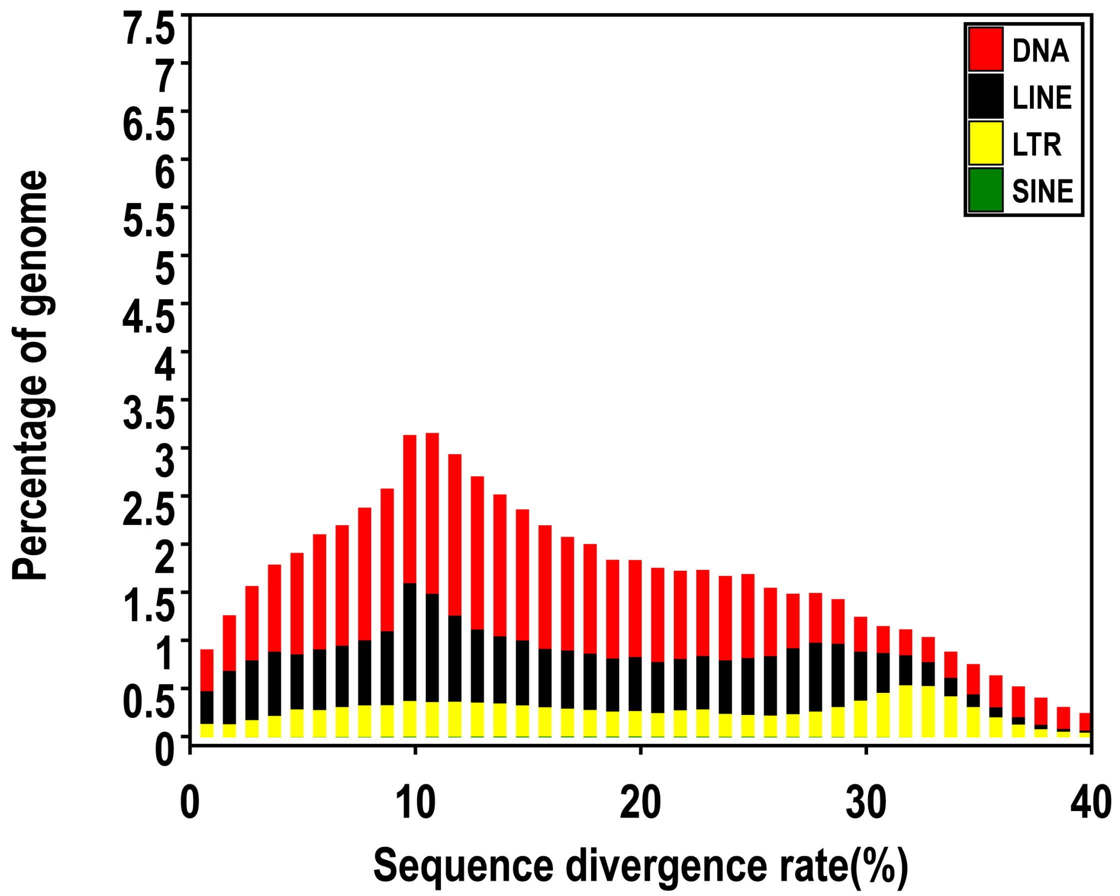


**Figure S2** Divergence rate of transposons in *Scylla paramamosain* genome

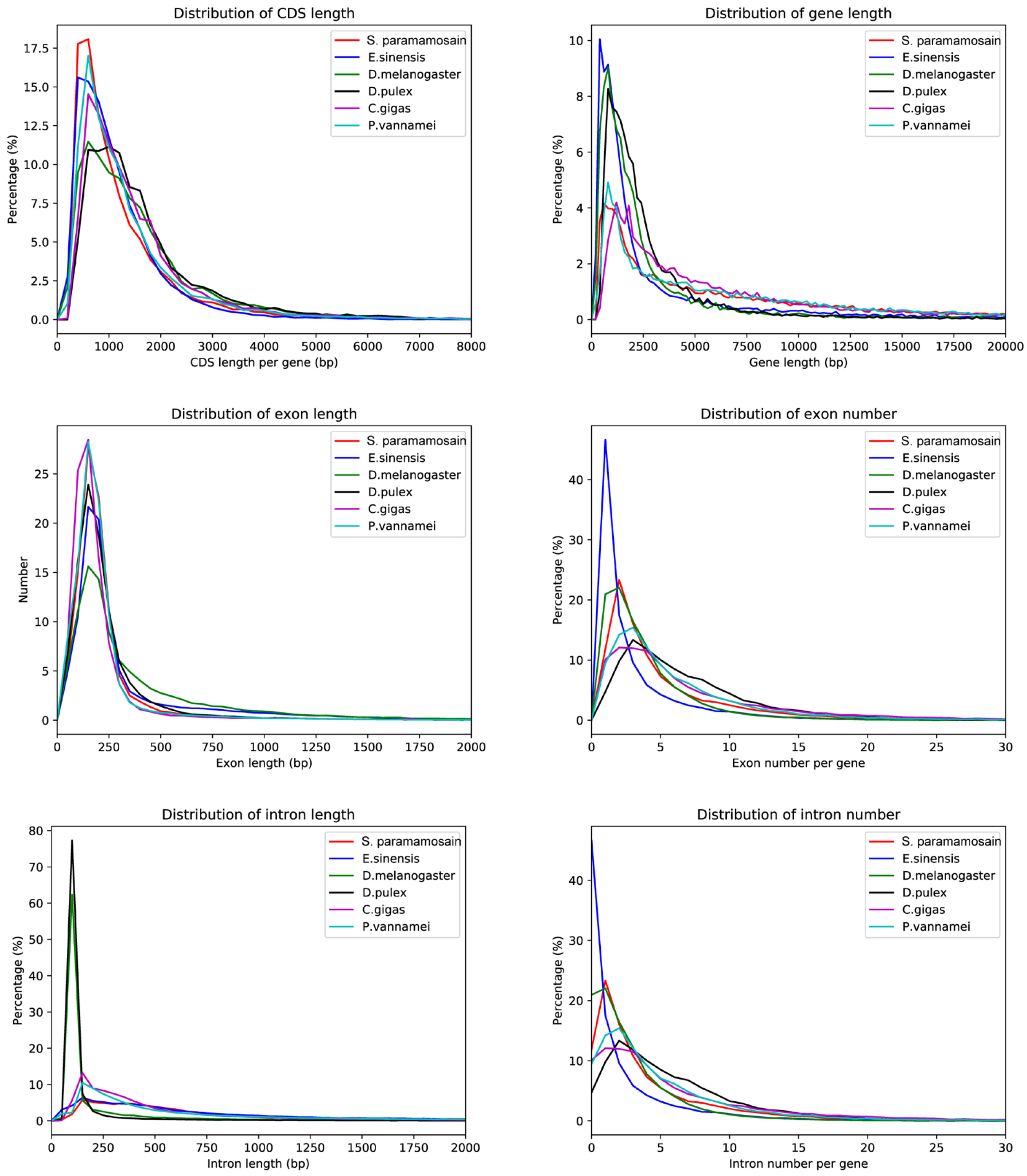


**Figure S3** Characterization of different genic regions in *Scylla paramamosain* and five other arthropods.

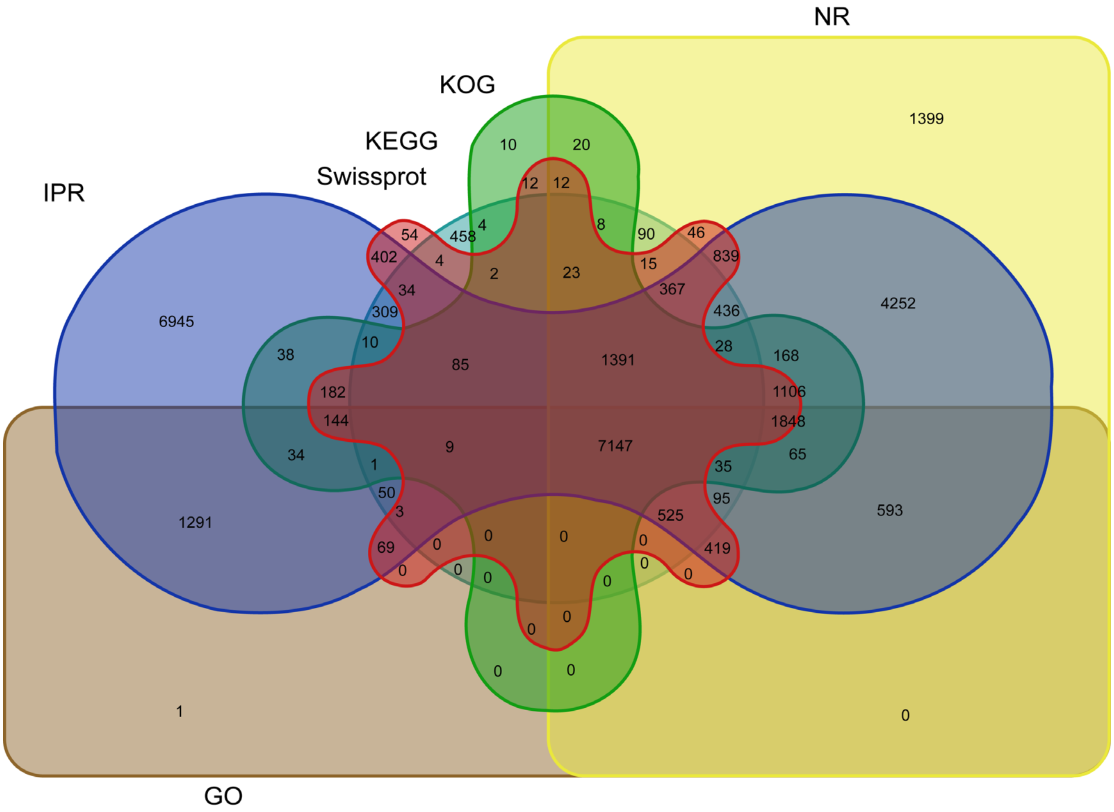


**Figure S4** Functional annotation of predicted genes in *Scylla paramamosain*

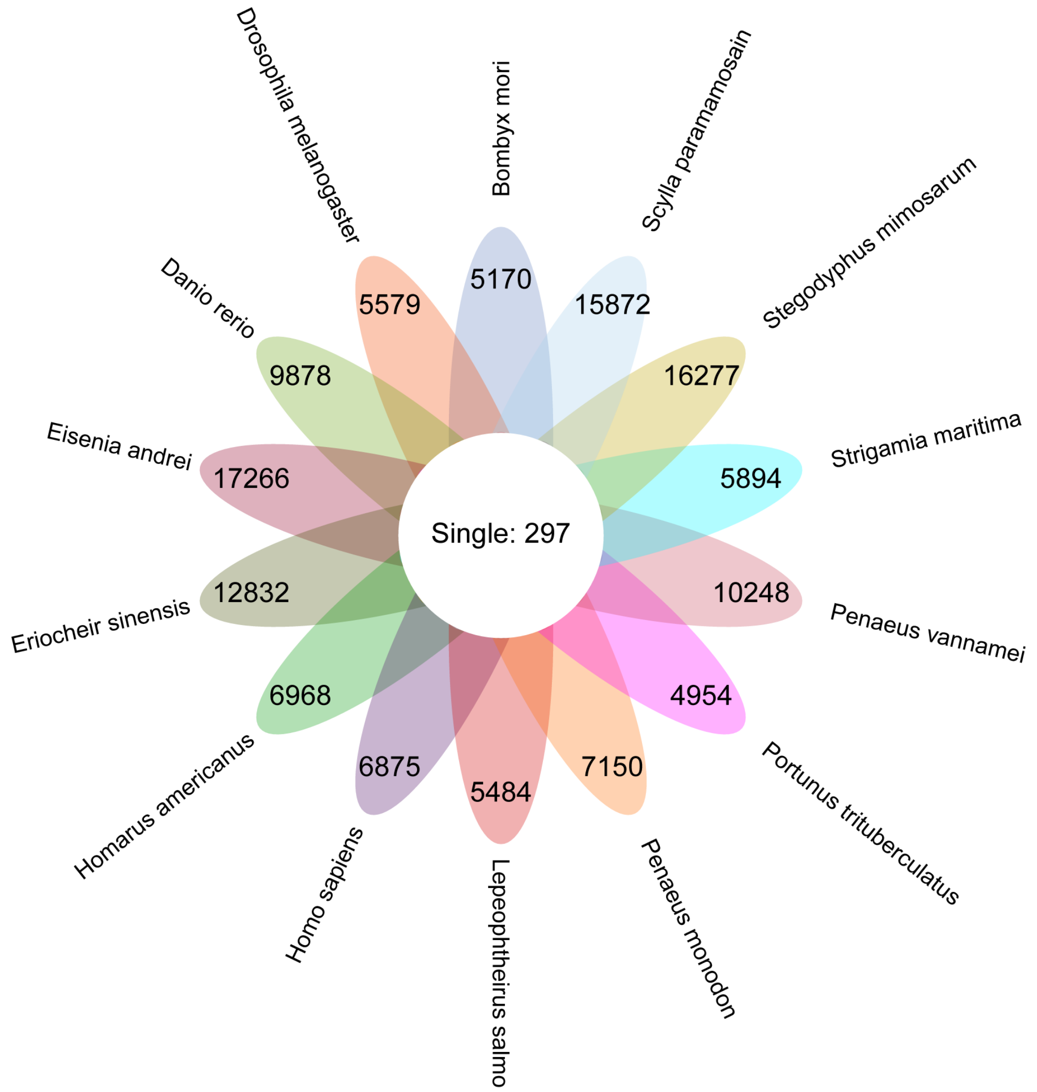


**Figure S5** Ortholog gene analysis and phylogenetic trees of 14 selected species

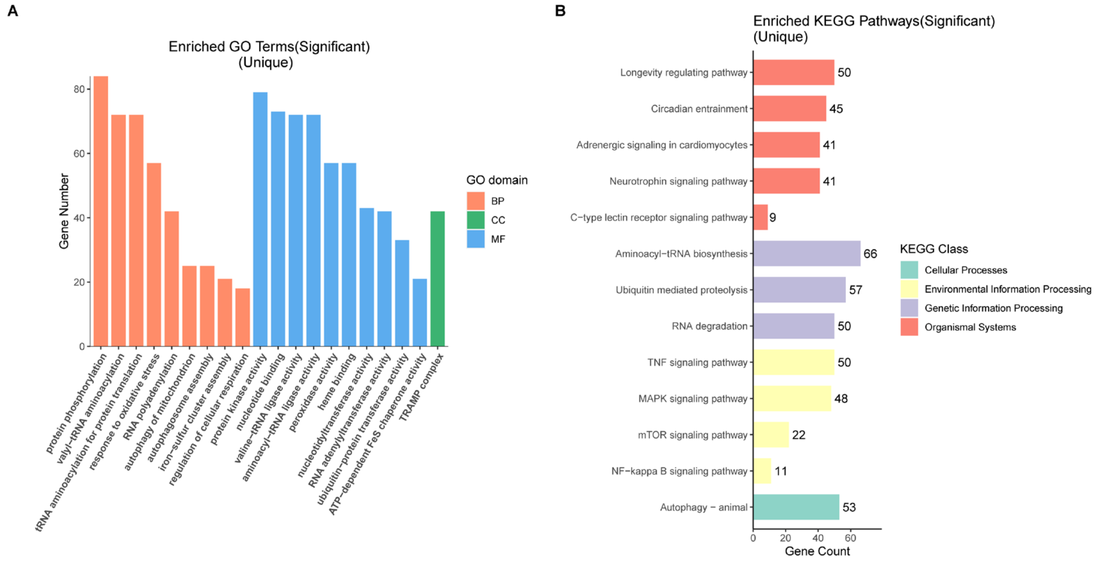


**Figure S6** GO (**A**) and KEGG (**B**) enrichment of unique genes in *Scylla paramamosain*

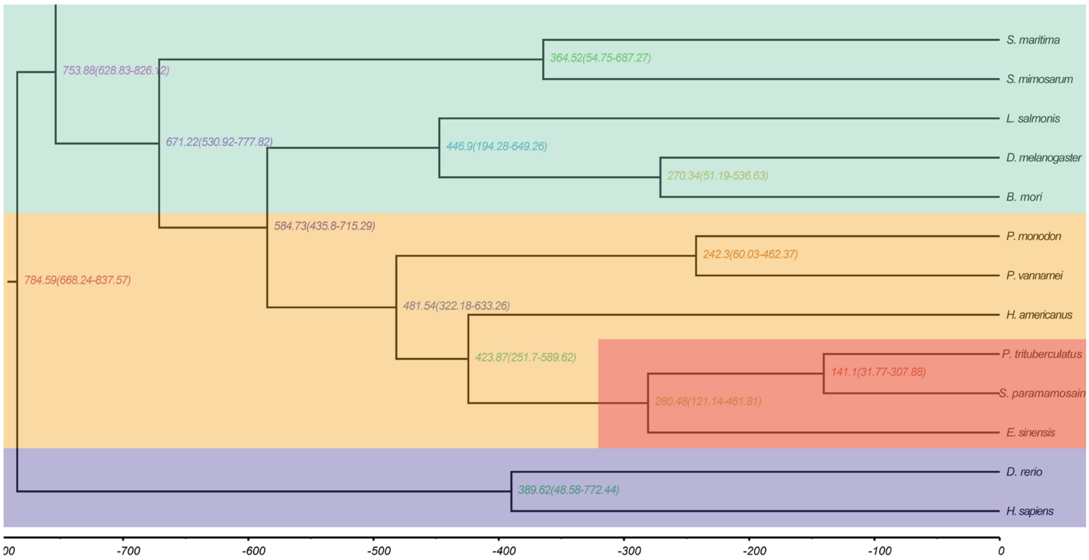


**Figure S7** Phylogenetic trees of 14 selected species including *Scylla paramamosain, Portunus trituberculatus, Eriocheir sinensis, Penaeus vannamei, Homarus americanus, Penaeus monodon, Drosophila melanogaster, Bombyx mori, Eisenia andrei, Lepeophtheirus salmonis, Stegodyphus mimosarum, Strigamia maritima, Danio rerio,* and *Homo sapiens*

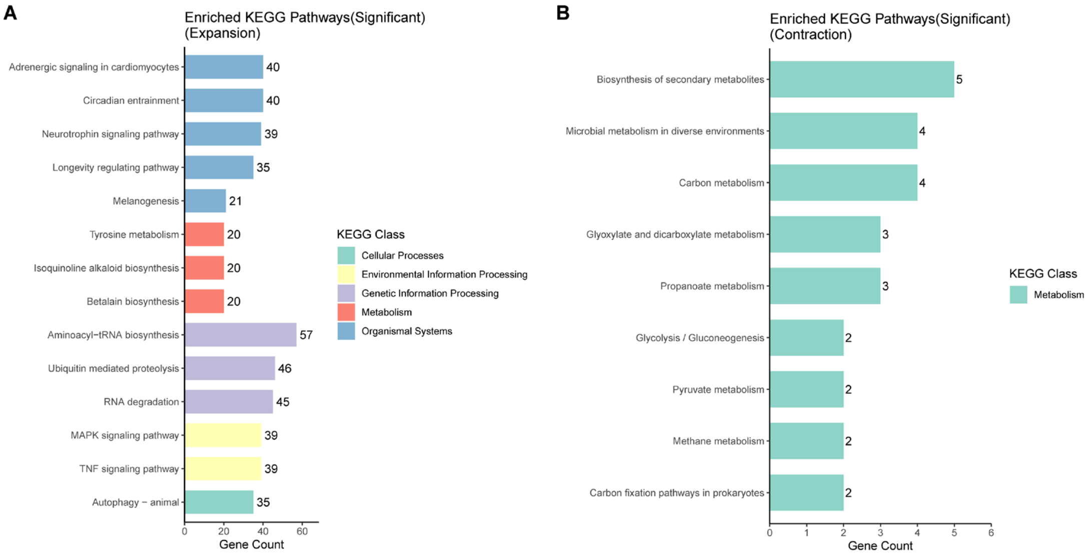


**Figure S8** KEGG enrichment of expansion (**A**) and contraction (**B**) genes in *Scylla paramamosain*

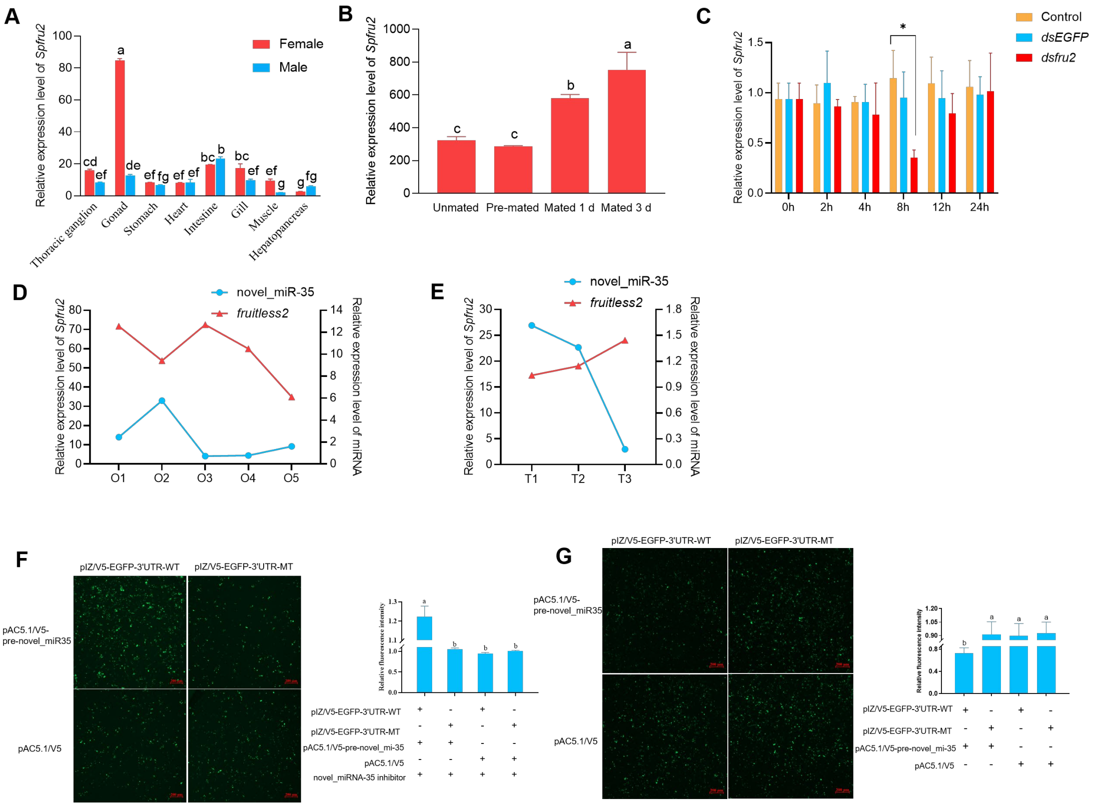


**Figure S9** The expression and regulation of fru2 genes in gonads of *Scylla paramamosain*. **(A)** Relative expression of *Spfru2* in different tissues of *Scylla paramamosain*. Bars with different lowercase letters indicate significant difference ($p < 0.05$). **(B)** Relative expressions of *SPfru2* in different mating stages of *Scylla paramamosain* female. **(C)** Expression changes of *Spfru2* in ovary after the interference of *fru2* expression. **(D)** Relative expression of novel-miR35 and *Spfru2* in different developmental stages of ovary. O1-O5: stage OI to stage OV. **(E)** Relative expression of novel-miR35 and *Spfru2* in different developmental stages of testis. T1-T3: stage TI to stage TIII. Bars with different lowercase letters indicate significant difference ($p < 0.05$). **(F)** The S2 cells were co-transfected with WT *fru2* 3′-UTR, and the mutated-type of *fru2* 3′-UTR (MT), together with novel-miR35 inhibitor and pre–novel_miR-35 plasmid. The data were expressed as the relative fluorescence intensity. **(G)** pre–novel_miR-35 plasmid or control plasmid were co-transfected WT *fru2* 3′-UTR, and the mutated-type of *fru2* 3′-UTR (MT) for 48 h. The data were expressed as the relative fluorescence intensity. $P < 0.05$.

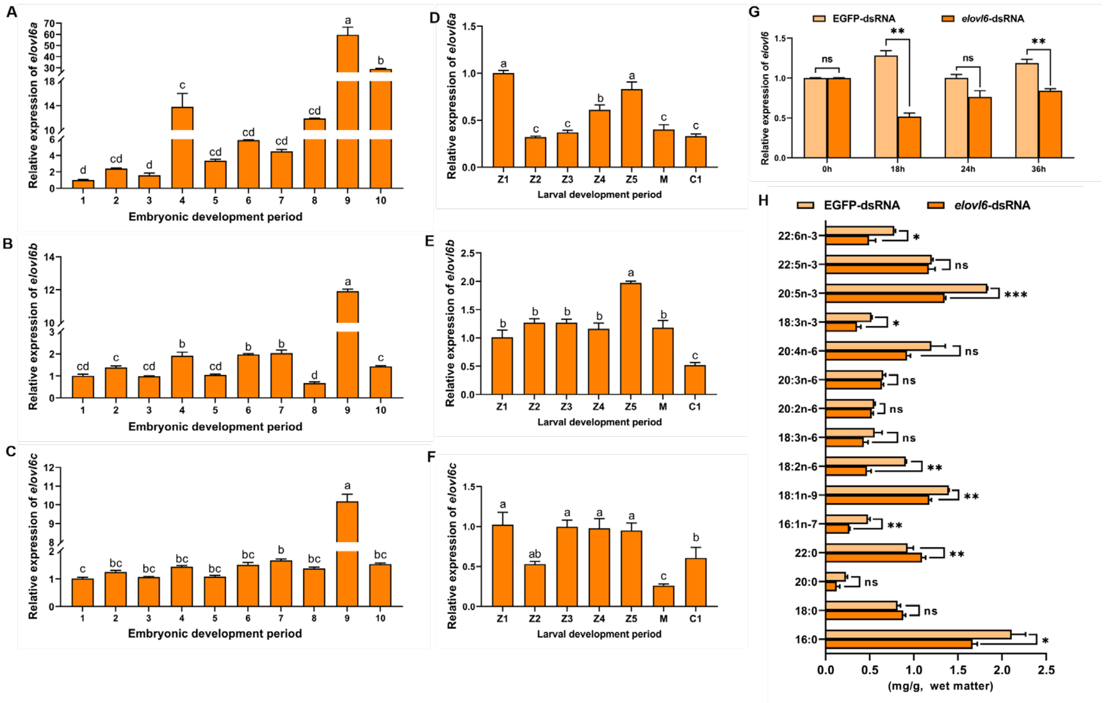


**Figure S10** Neo-functionalization of the *elovl6* gene in the LC-PUFA synthesis pathway in crustacean species. **(A)** Relative expressions of *elovl6a* in different developmental stages of embryo. 1: fertilized egg, 2: cleavage, 3: blastula, 4: gastrula, 5: nauplius, 6: five pairs of appendages, 7: seven pairs of appendages, 8: eye-pigment formation, 9: prehatching, 10: hatching. **(B)** Relative expressions of *elovl6b* in different developmental stages of embryo. **(C)** Relative expressions of *elovl6c* in different developmental stages of embryo. **(D)** Relative expressions of *elovl6a* in different developmental stages of larvae. **(E)** Relative expressions of *elovl6b* in different developmental stages of larvae. **(F)** Relative expressions of *elovl6c* in different developmental stages of larvae. **(G)** Expression levels of the *elovl6* gene after RNA interference in isolated hepatopancreas of green mud crab. **(H)** The contents of fatty acids in hepatopancreas after interfering the expression of elovl6 gene.

**Table S1 Comparison of published crustaceans genome**

| Species | Genome Size | Contig N50 | Scaffold N50 | Total number of genes | Average transcript length(bp) | Average CDS length(bp) | Average exons number per gene | Sequencing platform |
|---|---|---|---|---|---|---|---|---|
| *Scylla paramamosain* | 1.21 Gb | 11.45 Mb | 23.61 Mb | 33,662 | 22,869.83 | 1,187.71 | 6.06 | Nanopore, HiFi, HiC |
| *Eriocheir sinensis* | 1.57 Gb | 26.05 kb | 17.13 Mb | 28,033 | 4,578.54 | 1,074.94 | 3.24 | Pacbio, Illumina, HiC (Cui et al, 2021) |
| *Procambarus virginalis* | 3.29 Gb | 1.18 kb | 39.28 kb | 21,773 | 6,501.48 | 838.12 | 3.82 | Illumina, Hiseq 2000 (Gutekunst et al, 2018) |
| *Portunus trituberculatus* | 1.01 Gb | 4.12 Mb | 21.79 Mb | 16,796 | 11,195.08 | 1,465.56 | 6.6 | Nanopore, BGISEQ, HiC (Tang et al, 2020) |
| *Litopenaeus vannamei* | 1.66 Gb | 57 kb | 605 kb | 25,596 | 10,464.60 | 1,305.76 | 6.16 | Pacbio, Illumina, BAC (Zhang et al, 2019) |
| *Penaeus monodon* | 2.39 Gb | 79 kb | 44.86 Mb | 30,038 | 1,428 | | | Pacbio, Chicago, HiC (Uengwetwanit et al, 2021) |
| *Homarus americanus* | 2.29 Gb | 133.3 kb | 759.6 kb | 25,284 | 9,560 | 1,171 | 5.79 | Nanopore, Chicago, HiC (Polinski et al, 2021) |
| *Scylla paramamosain* | 1.55 Gb | 191 kb | 325 kb | 17,821 | | 1,527 | | Pacbio, Illumina, HiC (Zhao et al, 2021) |

**Table S2 Details of repeat sequences in *Scylla paramamosain***

| Class | Order | Super family | Number of elements | Length of sequence(bp) | Percentage of sequence(%) |
|---|---|---|---|---|---|
| Class I | | | 2158996 | 311984072 | 25.77 |
| | LINE | | 1164005 | 210695642 | 17.40 |
| | | Unknown | 112579 | 25034486 | 2.07 |
| | | CR1 | 221126 | 63321706 | 5.23 |
| | | CR1-Zenon | 115038 | 63628469 | 5.26 |
| | | L2 | 149069 | 10940876 | 0.90 |
| | | Penelope | 143776 | 8079613 | 0.67 |
| | | L1-Tx1 | 86092 | 3991730 | 0.33 |
| | | L1 | 113353 | 5277990 | 0.44 |
| | | R2-NeSL | 17665 | 1467079 | 0.12 |
| | | RTE-X | 20908 | 3640916 | 0.30 |
| | | R1 | 19475 | 1326564 | 0.11 |
| | | Rex-Babar | 22390 | 4570627 | 0.38 |
| | | RTE-BovB | 26669 | 6252341 | 0.52 |
| | | I-Jockey | 26536 | 2510007 | 0.21 |
| | | I | 10844 | 2450867 | 0.20 |
| | | Jockey | 3152 | 2132167 | 0.18 |
| | | Other | 75333 | 6070204 | 0.50 |
| | LTR | | 965998 | 99478958 | 8.22 |
| | | Gypsy | 328359 | 52422981 | 4.33 |
| | | ERV1 | 214468 | 7323356 | 0.60 |
| | | Unknown | 158516 | 23646993 | 1.95 |
| | | Pao | 42247 | 5584457 | 0.46 |
| | | ERVK | 132425 | 3968298 | 0.33 |
| | | Copia | 48946 | 2513227 | 0.21 |
| | | DIRS | 9729 | 2008485 | 0.17 |
| | | Other | 31308 | 2011161 | 0.17 |
| | SINE | | 28993 | 1809472 | 0.15 |

| | | | | | |
|---|---|---|---|---|---|
| Class II | | | 3006682 | 275205028 | 22.73 |
| | DNA | | 2881696 | 266918678 | 22.05 |
| | | P | 26684 | 1619787 | 0.13 |
| | | Unknown | 474614 | 86375737 | 7.13 |
| | | hAT-Ac | 321021 | 22462282 | 1.86 |
| | | Maverick | 220763 | 10879755 | 0.90 |
| | | hAT-Tip100 | 52202 | 2404302 | 0.20 |
| | | Ginger | 146399 | 7828469 | 0.65 |
| | | Kolobok-Hydra | 117795 | 10636249 | 0.88 |
| | | CMC-EnSpm | 395444 | 45344214 | 3.75 |
| | | PIF-Harbinger | 109827 | 5212437 | 0.43 |
| | | hAT-Blackjack | 47799 | 2134384 | 0.18 |
| | | Zisupton | 113881 | 7930998 | 0.66 |
| | | Kolobok-T2 | 46244 | 2426350 | 0.20 |
| | | Sola-1 | 83332 | 6221745 | 0.51 |
| | | PIF-Spy | 44858 | 2769776 | 0.23 |
| | | TcMar-Tc1 | 115337 | 4327601 | 0.36 |
| | | hAT-Charlie | 49113 | 2940755 | 0.24 |
| | | Sola-3 | 31175 | 1844837 | 0.15 |
| | | MULE-MuDR | 86652 | 9113921 | 0.75 |
| | | TcMar-Mariner | 38700 | 13984044 | 1.15 |
| | | TcMar-Fot1 | 42767 | 2510238 | 0.21 |
| | | CMC-Transib | 26917 | 2126293 | 0.18 |
| | | TcMar-Tigger | 10016 | 2460295 | 0.20 |
| | | PiggyBac | 10245 | 1235549 | 0.10 |
| | | Other | 269911 | 12128660 | 1.00 |
| | RC | | 111255 | 4050488 | 0.33 |
| | | Helitron | 111255 | 4050488 | 0.33 |
| | MITE | | 13731 | 4235862 | 0.35 |
| | | Unknown | 13731 | 4235862 | 0.35 |
| Total TEs | | | 5165678 | 587189100 | 48.50 |

**Table S3 Comparison of protein-coding genes between *Scylla paramamosain* and other species**

| Source | Number of gene | Average gene length(bp) | Average CDS length(bp) | Average exons per gene | Average exon length(bp) | Average intron per gene | Average intro length(bp) |
|---|---|---|---|---|---|---|---|
| *Scylla paramamosain* | 33,662 | 12,535.06 | 1,208.52 | 5.36 | 225.29 | 4.36 | 2,595.33 |
| *Eriocheir sinensis* | 27,832 | 4,578.54 | 1,074.94 | 3.24 | 331.41 | 2.24 | 1,561.62 |
| *Drosophila melanogaster* | 13,955 | 4,593.23 | 1,558.47 | 3.89 | 400.27 | 2.89 | 1,048.79 |
| *Daphnia pulex* | 15,295 | 3,393.58 | 1,630.89 | 6.91 | 235.89 | 5.91 | 298.06 |
| *Crassostrea gigas* | 28,402 | 7,404.91 | 1,473.73 | 7.54 | 195.51 | 6.54 | 907.22 |
| *Penaeus vannamei* | 24,974 | 10,464.60 | 1,305.76 | 6.16 | 212.06 | 5.16 | 1,775.78 |

**Table S4 Summary of non-coding genes in *Scylla paramamosain***

| Type | Number | Average_length(bp) | Total_length(bp) | Percentage(%) |
|---|---|---|---|---|
| regulatory | 361 | 47.94 | 17,307 | 0.0014 |
| tRNA | 3,153 | 73.85 | 232,845 | 0.0192 |
| ncRNA | 633 | 147.89 | 93,612 | 0.0077 |
| rRNA | 1,055 | 398.93 | 420,867 | 0.0348 |
| rRNA_stat: | | | | |
| 18S | 30 | 1,856.83 | 55,705 | 0.0046 |
| 28S | 34 | 7,286.85 | 247,753 | 0.0205 |
| 5S | 961 | 117.18 | 112,608 | 0.0093 |
| 5.8S | 30 | 160.03 | 4,801 | 0.0004 |
| ncRNA_stat: | | | | |
| other | 75 | 272.25 | 20,419 | 0.0017 |
| snRNA | 36 | 94.81 | 3,413 | 0.0003 |
| miRNA | 449 | 131.67 | 59,119 | 0.0049 |
| spliceosomal | 73 | 146.04 | 10,661 | 0.0009 |

**Table S5 Functional characterization of *Eriocheir sinensis, Portunus trituberculatus,* and *Scylla paramamosain* Elovl6 via heterologous expression in yeast *Saccharomyces cerevisiaea***

| FA | Substrate | Product | Conversion rate (%) | | | | | | | | | Activity |
|---|---|---|---|---|---|---|---|---|---|---|---|---|
| | | | *Eselovl6-X1* | *Eselovl6-X2* | *Eselovl6-X3* | *Ptelovl6-X1* | *Ptelovl6-X1* | *Ptelovl6-X1* | *Spelovl6a* | *Spelovl6b* | *Spelovl6c* | |
| SFA | 16:00 | 18:00 | 4.34±0.22 | 8.04±0.03 | 18.12±0.41 | 4.56±0.25 | 8.60±0.29 | 34.11±0.79 | 31.13±0.83 | 9.11±0.43 | 11.89±0.68 | C16→C18 |
| | | 20:00 | 1.86±0.21 | 1.91±0.17 | 1.71±0.11 | 1.97±0.30 | 2.42±0.10 | 3.10±0.22 | 2.35±0.18 | 2.49±0.13 | 2.96±0.05 | C18→C20 |
| | | 22:00 | 33.01±0.94 | 36.50±0.49 | 39.63±0.68 | 27.39±0.59 | 35.48±0.30 | 37.21±0.62 | 28.61±0.92 | 10.67±0.74 | 21.71±0.46 | C20→C22 |
| MUFA | 16:1n-7 | 18:1n-7 | 2.78±0.17 | 4.80±0.23 | 10.85±0.24 | 2.75±0.15 | 5.18±0.24 | 13.71±0.42 | 7.60±0.23 | 2.51±0.11 | 4.46±0.23 | C16→C18 |
| | 18:1n-9 | 20:1n-9 | 0.31±0.03 | 1.71±0.07 | 2.63±0.07 | 0.33±0.02 | 2.36±0.15 | 4.86±0.14 | 5.30±0.40 | 0.37±0.04 | 4.83±0.27 | C18→C20 |
| | | 22:1n-9 | 58.43±1.12 | 27.67±0.52 | 23.41±0.69 | 63.30±0.83 | 26.90±0.50 | 22.47±0.75 | 20.83±0.91 | 77.33±1.54 | 36.51±0.56 | C20→C22 |
| PUFA | 18:2n-6 | 20:2n-6 | 0.32±0.06 | 2.67±0.12 | 3.98±0.16 | 0.43±0.05 | 2.68±0.12 | 6.57±0.16 | 6.24±0.15 | 0.46±0.09 | 3.46±0.23 | C18→C20 |
| | 18:3n-6 | 20:3n-6 | 0.35±0.05 | 2.72±0.20 | 5.94±0.17 | 0.48±0.09 | 7.80±0.18 | 8.07±0.20 | 6.29±0.20 | 0.78±0.10 | 4.28±0.19 | C18→C20 |
| | 18:3n-3 | 20:3n-3 | 0.36±0.06 | 3.40±0.29 | 8.23±0.31 | 0.61±0.08 | 5.80±0.23 | 8.47±0.30 | 14.51±0.33 | 0.79±0.06 | 4.82±0.36 | C18→C20 |
| | 18:4n-3 | 20:4n-3 | 0.37±0.06 | 1.13±0.09 | 3.91±0.23 | 0.59±0.10 | 2.61±0.21 | 4.52±0.20 | 8.13±0.18 | 0.45±0.07 | 2.89±0.23 | C18→C20 |
| | 20:4n-6 | 22:4n-6 | ND | ND | ND | ND | ND | ND | ND | ND | ND | C20→C22 |
| | 20:5n-3 | 22:5n-3 | ND | ND | ND | ND | ND | ND | ND | ND | ND | C20→C22 |

**Table S6 Primers used for Eselovl6, Ptelovl6, and Spelovl6 functional characterization**

| Transcript | Primer | Sequence |
|---|---|---|
| *Eselovl6* | *Eselovl6*-X1-F | GAATATTAAGCTTGGTACCGAGCTCGGATCCATGACCGCAATGGACGCCTTGGCCAAGAGGTCCTTCTTCGACAACACGA |
| | *Eselovl6*-X1-R | GCCGCCAGTGTGATGGATATCTGCAGAATTCTTATTCCAGTTTACCCTTGGTGCTCTTTCCTTCGTAGGCGAGAGACTCCTGG |
| | *Eselovl6*-X2-F | GAATATTAAGCTTGGTACCGAGCTCGGATCCATGGAGAGCGAAAAGATGTTCTTCATCAATTTGACGAACGTGGAGGC |
| | *Eselovl6*-X2-R | GCCGCCAGTGTGATGGATATCTGCAGAATTCTTATTCCAGTTTACCCTTGGTGCTCTTTCCTTCGTAGGCGAGAGA |
| | *Eselovl6*-X3-F | GAATATTAAGCTTGGTACCGAGCTCGGATCCATGGAGTCGGTCACCATGCCTAACTATTCTTACGTCTTC |
| | *Eselovl6*-X3-R | GCCGCCAGTGTGATGGATATCTGCAGAATTCTTATTCCAGTTTACCCTTGGTGCTCTTTCCTTCGTAGGCGAGAGA |
| *Ptelovl6* | *Ptelovl6*-X1-F | GAATATTAAGCTTGGTACCGAGCTCGGATCCATGACAAGGAGCATGGAGACACTGCACAAATCTAGCTTCTTTGAAAACA |
| | *Ptelovl6*-X1-R | GCCGCCAGTGTGATGGATATCTGCAGAATTCTTATTCCAGTTTACCCTTGCTACCTTTTCCTTCATAAGCAATAGACTCCTT |
| | *Ptelovl6*-X2-F | GAATATTAAGCTTGGTACCGAGCTCGGATCCATGAGTTTCTTCATCAATTTGACGAACGTGGAGGCAGCTG |
| | *Ptelovl6*-X2-R | GCCGCCAGTGTGATGGATATCTGCAGAATTCTTATTCCAGTTTACCCTTGCTACCTTTTCCTTCATAAGCAATAGACT |
| | *Ptelovl6*-X3-F | GAATATTAAGCTTGGTACCGAGCTCGGATCCATGGAGTCGGTCACCATGCCAAATTATTCCTATGTCTTCAAGTTCGAGG |
| | *Ptelovl6*-X3-R | GCCGCCAGTGTGATGGATATCTGCAGAATTCTTATTCCAGTTTACCCTTGCTACCTTTTCCTTCATAAGCAATAGACTCCTTG |
| *Spelovl6* | *Spelovl6a*-F | GAATATTAAGCTTGGTACCGAGCTCGGATCCATGGAGTCGGTCACCATGCCAAATTATTCCTATG |
| | *Spelovl6a*-R | CCGCCAGTGTGATGGATATCTGCAGAATTCTTATTCCAGTTTACCCTTGCTACCTTTCCCTTCATAA |
| | *Spelovl6b*-F | GAATATTAAGCTTGGTACCGAGCTCGGATCCATGACAAGGAATATGGAGACACTGCATAAAT |

| | |
|---|---|
| *Spelovl6b*-R | CCGCCAGTGTGATGGATATCTGCAGAATTCTTATTCCAGTTTACCCTTGCTACCTTTCCCTTC |
| *Spelovl6c*-F | GAATATTAAGCTTGGTACCGAGCTCGGATCCATGAGTTTCTTCATCAATTTGACGAACGTGGAGGC |
| *Spelovl6c*-R | CCGCCAGTGTGATGGATATCTGCAGAATTCTTATTCCAGTTTACCCTTGCTACCTTTCCCTT |
